# Unifying the Electron Microscopy Multiverse through a Large-scale Foundation Model

**DOI:** 10.1101/2025.04.13.648639

**Authors:** Liuyuan He, Ruohua Shi, Wenyao Wang, Guanchen Fang, Yu Cai, Lei Ma

## Abstract

Accurate analysis of electron microscopy (EM) images is essential for exploring nanoscale biological structures, yet data heterogeneity and fragmented workflows hinder scalable insights. Pretrained on large, diverse datasets, image foundation models provide a robust framework for learning transferable representations across tasks. Here, we introduce EM-DINO, the first EM image foundational model pretrained on EM-5M, a large curated and standardized EM corpus (5 million images) encompassing multiple species, tissues, protocols, and resolutions. EM-DINO’s multi-scale embeddings capture rich image features that support multiple applications, including organ-specific pattern recognition, image deduplication, and high quality image restoration. Building on these representations, we developed OmniEM, a U-shaped architecture for unified dense prediction that achieves superior performance compared with task-specific models in both image restoration and segmentation. In restoration benchmarks, OmniEM matches the performance of the EM-specific diffusion model while reducing spurious structural artifacts that could mislead interpretation. It also outperforms previous methods across 2D and 3D mitochondrial segmentation, as well as multi-class organelle segmentation tasks. Furthermore, we demonstrate OmniEM’s integrated capability to generate high-resolution segmentations from low-resolution inputs, offering the potential to enable fine-scale subcellular analysis in legacy and high-throughput EM datasets. Together, EM-5M, EM-DINO, OmniEM, and an integrated Napari plugin comprise a comprehensive end-to-end toolkit for standardized EM analysis, advancing cellular and subcellular understanding and accelerating the discovery of novel organelle morphologies and disease-related alterations.

## Main

Electron microscopy (EM) provides an unparalleled window into the nanometer-scale architecture of biological systems, revealing subcellular organelles, synaptic connections, and tissue ultrastructure with extraordinary detail^1–6^. In recent years, high-throughput EM has transformed our ability to visualize cellular and tissue organization at this resolution. While traditional 2D EM relies on expert interpretation of densely packed organelles, volumetric EM (vEM) techniques have emerged over the past two decades to satisfy connectomics’ demand for large-scale neuronal reconstructions—generating datasets far too extensive for manual analysis or hypothesis testing^7–9^. Consequently, automated methods are indispensable in EM workflows, spanning low-level tasks (restoration, alignment), segmentation, and higher-order, question-driven analyses such as mapping synaptic connectivity at the circuit level. Unlike fluorescence microscopy, where semantic content derives from labels or contrast agents, EM images are grayscale and rich with nanometer-scale structural information. Therefore, EM research demands bespoke computational models and automated tools to fully harness modern EM’s potential and address the pressing biological questions it can illuminate^10^.

Modern deep-learning (DL) methods have driven substantial progress in EM image analysis, automating tasks such as segmentation, denoising, and registration^11–16^. Task-specific models now achieve expert-level performance on benchmarks For example, neuronal reconstruction (SNEMI3D^17^, CREMI^18^), organelle identification (MitoNet^13^, BetaSeg^19^), and image restoration (EMDiffuse^15^). However, these approaches depend on narrowly defined training sets and often degrade when applied to new imaging conditions or biological contexts^10^. Moreover, EM workflows inherently require distinct models at each processing stage (restoration, alignment, segmentation, and higher-level analysis) imposing significant overhead for tool integration and adaptation^1,20,21^, and confining DL solutions to limited task domains (e.g., specialized connectomics pipelines^22^). As a result, existing DL strategies lack the general-purpose flexibility needed for scalable, reproducible EM analysis.

Self-supervised image foundation models (FMs) offer a promising path toward unified, scalable EM workflows. By pretraining on large, diverse unlabeled datasets, FMs learn rich, transferable representations that can be adapted to a wide range of downstream tasks with minimal annotation^23^. In medical imaging, FMs have already shown notable success: pretrained models can be fine-tuned for segmentation, classification, and detection across heterogeneous datasets^24–26^. In EM, however, recent efforts have focused primarily on prompt-based adaptations of the Segment Anything Model (SAM) such as microSAM for general microscopy^27^, DendriteSAM for neuronal processes^28^, and TriSAM for vascular tracing^29^. These approaches depend on user-provided prompts (e.g., bounding boxes, masks), limiting scalability across the full spectrum of EM tasks from low-level restoration to high-level semantic analysis. Meanwhile, generative models have been applied to restoration^15^ but do not learn universally transferable representations. Thus, the EM field still lacks a unified, representation-centric foundation model framework capable of supporting robust, flexible analysis across diverse biological and technical conditions.

To address these gaps, we introduce a modular EM foundation pipeline comprising:

- **EM-5M**: A deduplicated, stratified corpus of 5 million 2D EM images drawn from 500 TB of raw data (EMPAIR^30^,
- **EM-DINO**: A vision transformer pretrained on EM-5M via the DINOv2 self-distillation framework^31^, producing robust, multi-scale embeddings that capture ultrastructural features.
- **OmniEM**: A U-shaped hybrid network that consumes EM-DINO’s final and intermediate patch embeddings and supports universal dense prediction EM tasks. OmniEM achieves strong and competitive performance in denoising, super-resolution, single-organelle, and multi-organelle segmentation, with native support for both 2D slice-based inputs and 3D volumetric EM data.
- **OmniEM-Napari**: An interactive Napari plugin that packages pretrained EM-DINO and OmniEM checkpoints into a user-friendly GUI, enabling researchers (including those without extensive coding expertise) to deploy OmniEM workflows on their own EM data.

Figure 1 offers a schematic overview of our workflow: large-scale data curation, self-supervised pretraining, and downstream task adaptation. Although foundation-model strategies are well established in natural and medical imaging, EM data pose unique challenges that vEM produces many nearly identical, contiguous slices (< 70 nm apart), so naively including adjacent patches would introduce data redundancy and imbalance, degrading dataset quality^32^. To mitigate this, we devised a bespoke deduplication pipeline that clusters patches by EM-DINO embeddings or spatial proximity, followed by uninformative filtering (Extended Data Table 1). The data cleaning and model training loop, alternating between dataset refinement and EM-DINO checkpoint updates, yields a highly diverse, nonredundant EM-5M corpus and a robust foundational model(Fig. 1, Left).

**Fig. 1.**
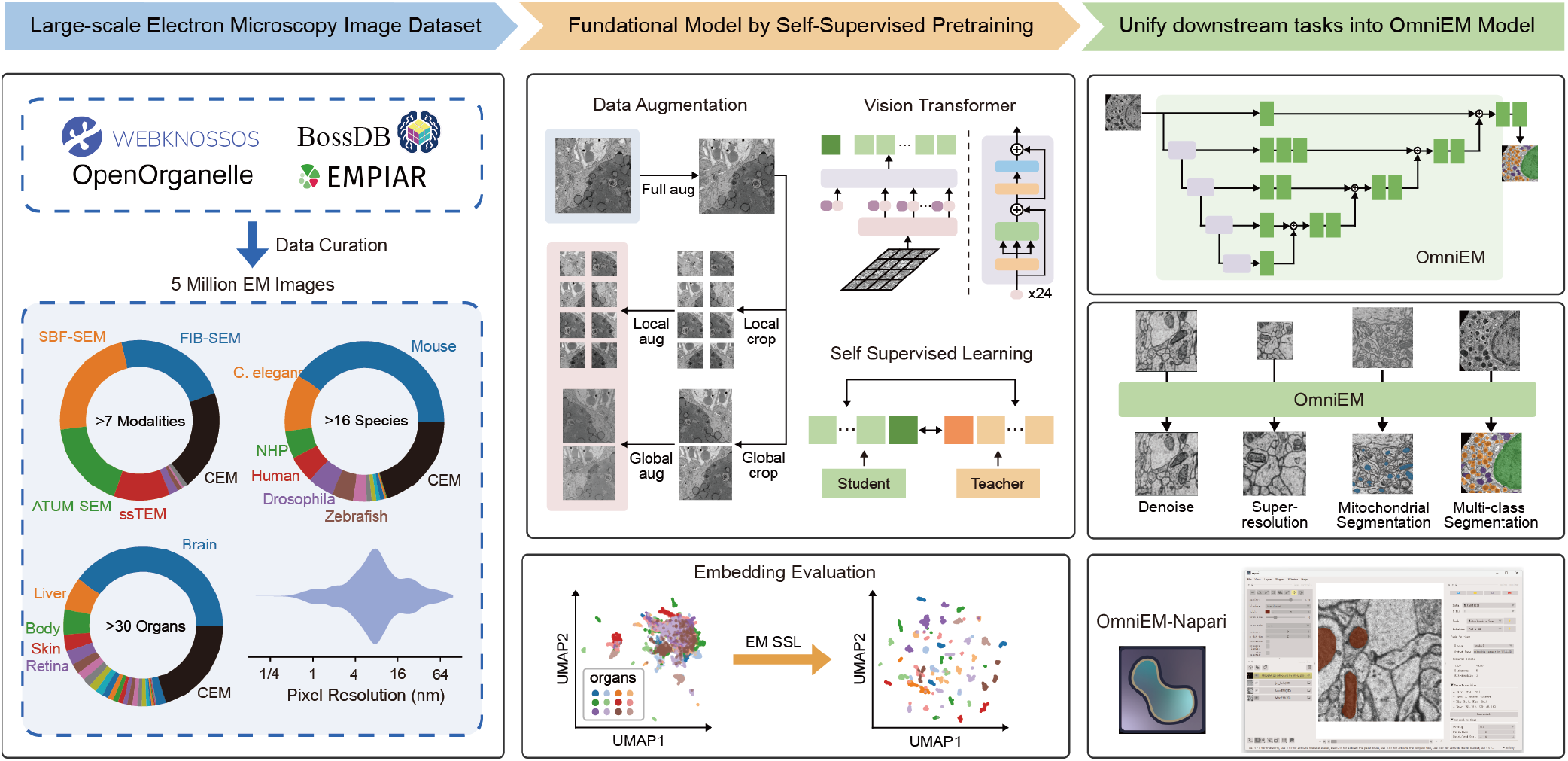
Overview of *EM-5M, EM-DINO*, and *OmniEM*. **Left**. Data from Webknossos, BossDB, OpenOrganelle, EMPIAR and other sources are collected and curated to *EM-5M* through a unified preprocessing pipeline, including standardization, patch extraction, deduplication and uninformative-region filtering. EM-5M contains over 5 million curated andstandardized images, spanning more than seven imaging modalities, over 16 animal species, more than 30 organ types, and a broad range of pixel sizes (0.5–70 nm). **Middle**. Self-supervised pretraining of *EM-DINO* on *EM-5M* using EM-specific data augmentations to learn transferable multi-scale embeddings for EM images **Right**. From top to bottom: the OmniEM architecture that consumes intermediate and final EM-DINO embeddings for dense prediction, representative downstream tasks (image restoration and segmentation), and the interface of the Napari pluginfor interactive deployment.

Next, we use self-supervised learning to pretrain EM-DINO on EM-5M (Figure 1, Middle Up). In organ classification benchmarks, EM-DINO embeddings formed distinct clusters reflecting organ-specific morphology (Figure 1, Middle Bottom), confirming that large, heterogeneous EM data produce transferable, multi-scale representations. In principle, the EM-DINO embedding space can serve as a unified representation supporting low-level processing (alignment, denoising, super-resolution), mid-level understanding (cell and organelle segmentation), and high-level analysis (connectomics, spatial organization of subcellular structures).

To operationalize this potential, we developed OmniEM, which translates EM-DINO embeddings into dense predictions across multiple tasks (Figure 1, Right Up). OmniEM’s architecture extracts patch-level features from EM-DINO’s final and intermediate layers, upsamples them via a multi-scale decoder, and applies task-specific heads for restoration or segmentation. We demonstrate OmniEM’s utility in denoising, super-resolution, mitochondrial segmentation, and multi-class segmentation (Figure 1, Right Middle). Finally, we integrated pretrained OmniEM checkpoints into Napari-OmniEM for seamless deployment (Figure 1, Right Bottom).

Overall, this work proposes a pipeline for EM image representation learning that combines large-scale self-supervised learning with EM-specific data curation. We publicly release the full toolkit, including EM-5M, EM-DINO, OmniEM, as well as data curation scripts, along with an online service for deploying pretrained segmentation models. These resources support a wide range of community use, from out-of-the-box applications to developing new downstream tasks and expanding curated EM datasets. In the sections that follow, we first evaluate EM-DINO embeddings via organ classification benchmarks (Figure 2). We then assess OmniEM’s performance on key downstream tasks—image restoration (Figure 3) and both mitochondrial and multi-organelle segmentation (Figure 4). Lastly, we present practical applications, illustrating how OmniEM’s modules can be combined in compound workflows and deployed via the Napari plugin (Figure 5). Collectively, these results highlight the power of EM foundation models to enable scalable, reproducible, end-to-end automation and user-guided refinement in EM analysis. Given the diversity of EM imaging modalities, we anticipate this study will provide a strong reference foundation for more general-purpose approaches to EM image understanding, with further improvements expected as larger datasets become available.^1^

**Fig. 2.**
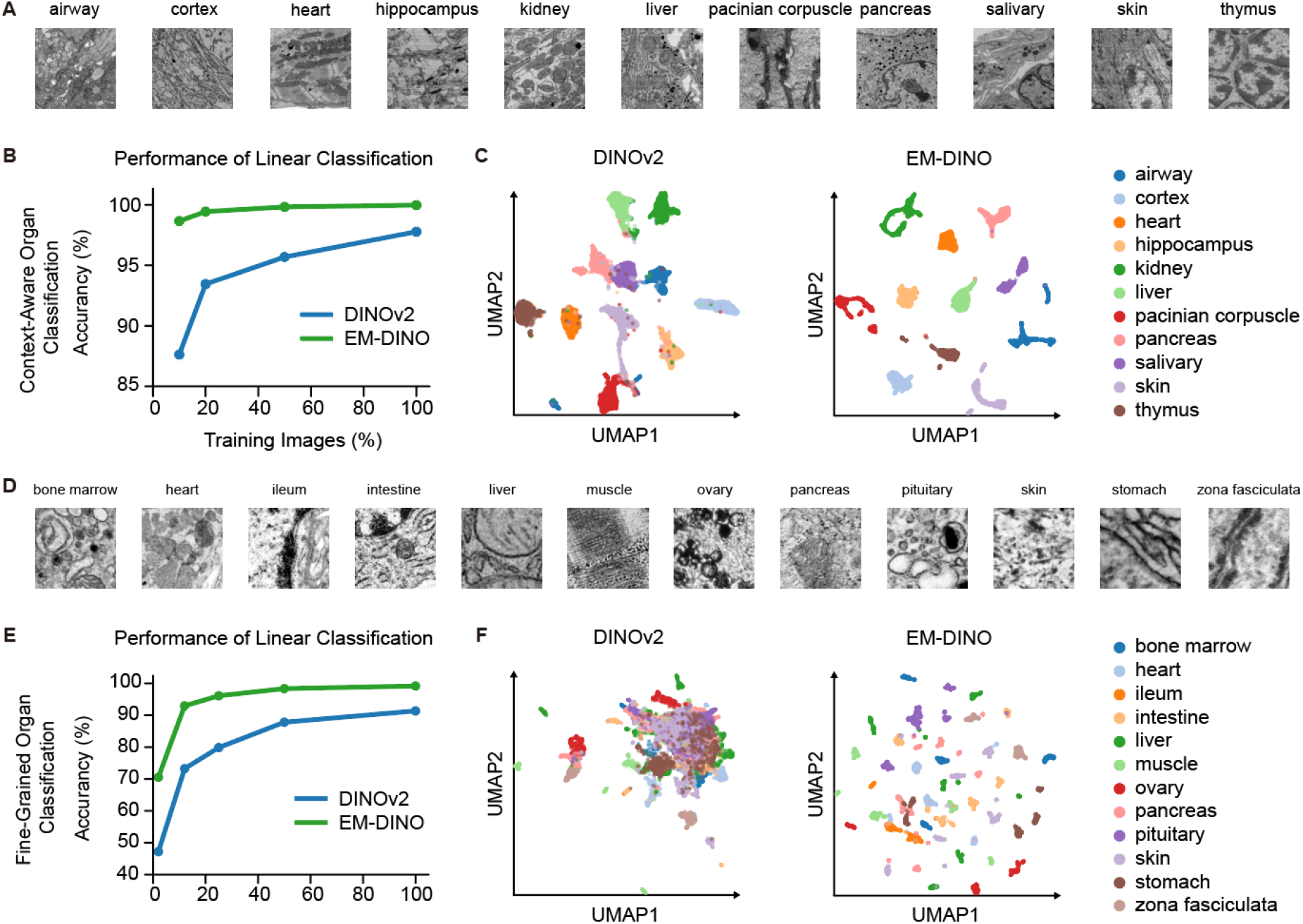
Evaluation of EM-DINO embeddings on organ classification tasks. *EM-DINO*, pretrained on *EM-5M*, is compared with a natural image–pretrained *DINOv2* backbone under the same downstream evaluation protocol. (**A**) Schematic of the context-aware classification task using morphologically rich EM image patches from 11 organ types. (**B**) Classification accuracy of *DINOv2* and *EM-DINO* across varying amounts of training data. (**C**) UMAP visualization of frozen image embeddings from the context-aware task, showing clearer organ-level separation with *EM-DINO*. (**D**) Schematic of the fine-grained classification task using small field-of-view, low-context EM images from 12 organs in an independent dataset not used during pretraining. (**E**) Classification accuracy of both models on the fine-grained task across different training data sizes. (**F**) UMAP visualization of frozen image embeddings in the fine-grained setting, where *EM-DINO* forms more distinct and semantically meaningful clusters.

**Fig. 3.**
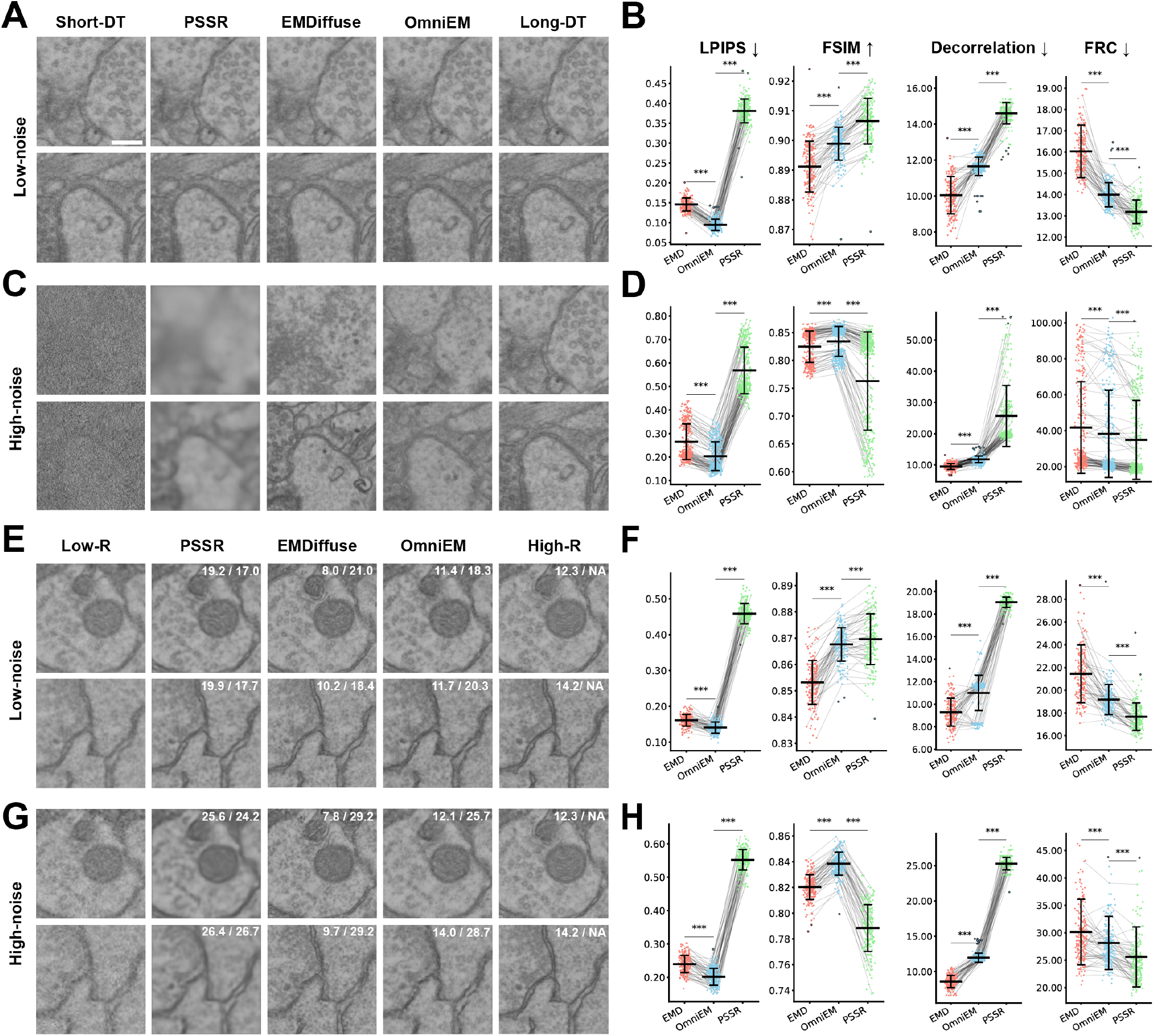
Evaluation of OmniEM image denoising and super-resolution. **(A)** Comparison of the denoising results from different models under low noise conditions. From left to right: the noisy image acquired with short dwell time (used as model input), the restored results from PSSR, EMDiffuse, and OmniEM, and the high-quality image acquired with long dwell time (used as ground truth reference). Two images are shown as representative cases. **(B)** Graphs comparing LPIPS, FSIM, and image resolution (assessed using decorrelation and Fourier ring correlation (FRC)) across different models. LPIPS, FSIM, and FRC resolution values were computed between each restored image and its corresponding ground truth, whereas decorrelation-based resolution was estimated independently of ground truth (n = 191 image tiles; xy dimensions, 224 × 224 pixels). Arrows next to each metric indicate the direction of better performance. Values are shown as mean ± s.d. P values are indicated for comparisons between OmniEM and the other two models (*P < 0.05, **P < 0.01, ***P < 0.001; NS, not significant; paired t-test). Lines connecting scatter points denote paired measurements from the same image tile restored by different models. For visualization clarity, only 25% of data points are displayed. Data points exceeding mean ± 3 × s.d. were considered outliers, highlighted with black outlines, and excluded from statistical analysis. **(C and D)** Shown are the corresponding qualitative and quantitative denoising results under high noise conditions, with the same layout and legend conventions as in A and B (n = 316 image tiles), respectively. **(E)** Comparison of the super-resolution results from different models under low noise conditions. From left to right: the low-resolution input image, super-resolved outputs from PSSR, EMDiffuse, and OmniEM, and the high-resolution ground truth image. Two representative images are shown. Resolution values estimated using decorrelation and FRC are indicated in the upper right corner of each restored image. **(F)** Shown are the paired performance comparisons between different models under low noise conditions for super-resolution, using the same four metrics. Layout and legend conventions follow those in B and D (n = 179 image tiles). **(G and H)** Shown are the corresponding qualitative and quantitative super-resolution results under high noise conditions, following the same layout and conventions as in E and F (n = 189 image tiles). Scale bar: 200 nm.

**Fig. 4.**
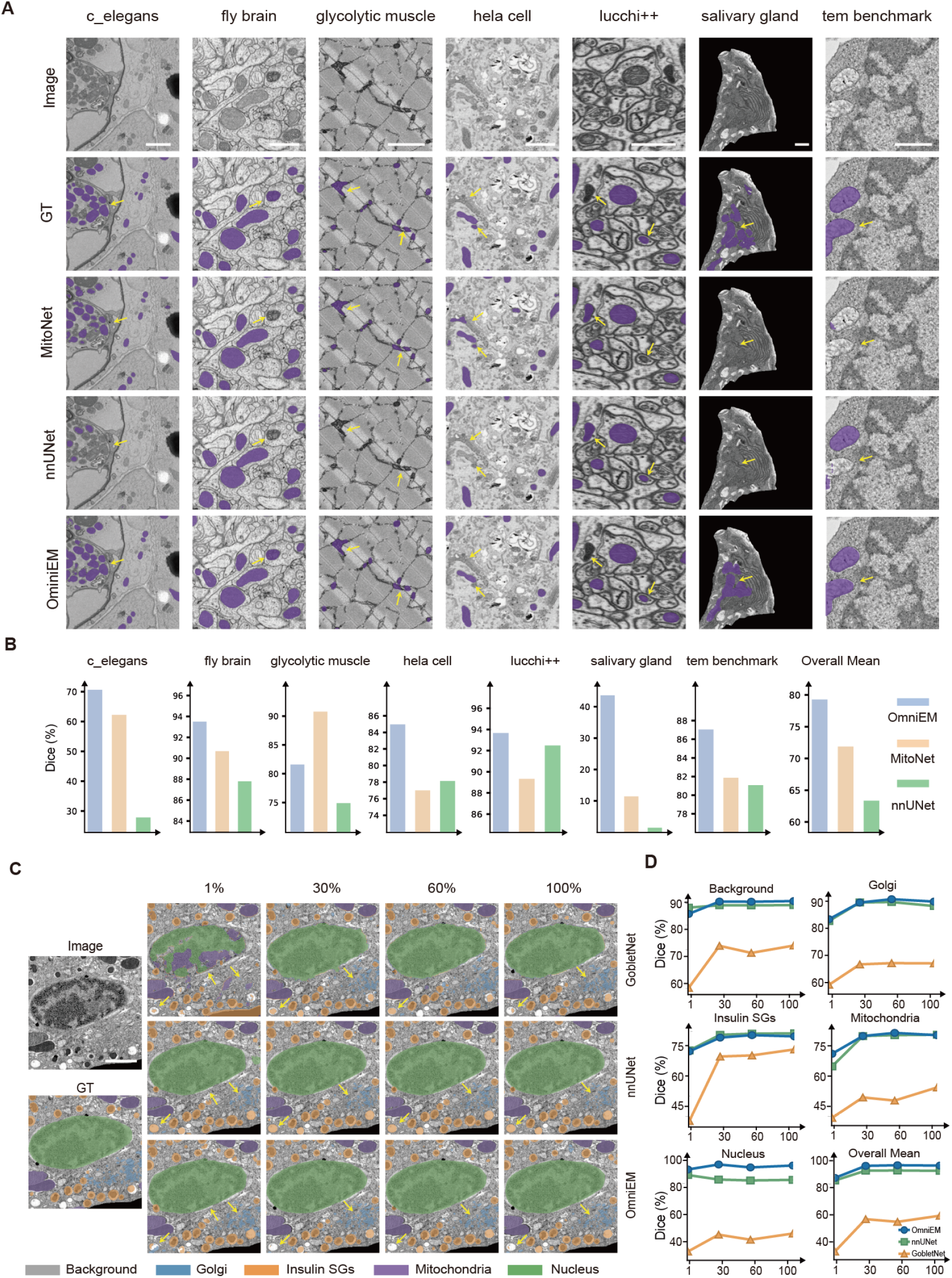
Segmentation for mitochondria and multi-organelle. (**A**) Seven test datasets from CEM-MitoLab mitochondrial segmentation benchmark, from left to right: C. elegans, fly brain, glycolytic muscle, HeLa cell, Lucchi++, salivary gland, and TEM images benchmark. From top to bottom: raw input, ground truth (GT), MitoNet predictions, nnU-Net predictions and OmniEM predictions. Purple masks represent GT labels or predicted segmentations overlaid on the raw input images. Yellow arrows highlight regions where OmniEM delineates mitochondrial boundaries more accurately, whereas MitoNet or nnU-Net tends to over-segment or miss these structures. **(B)** Dice coefficient comparisons between OmniEM (light blue) MitoNet (light apricot), and nnU-Net (green) across the corresponding seven benchmark datasets in A; the final inset summarizes the mean Dice scores across all datasets. (**C**) Representative EM images and segmentation results on Betaseg dataset (2D experiment). Column one, raw input images and GT labeling. Columns two to five, segmentation results at the training ratio of 1%, 30%, 60%, and 100%, respectively, with results displayed from top to bottom for GobletNet, nnU-Net, and OmniEM. Yellow arrows indicate regions where OmniEM precisely segments while GobletNet or nnU-Net over-segments or misses. Colors: background, grey; Golgi apparatus, blue; insulin SGs, orange; mitochondria, purple; nucleus, green. (**D**) Class-specific Dice coefficient curves for OmniEM (blue) nnU-Net (green), and GobletNet (orange) across training ratios. Scale bar, 1 *µ*m, applies to all insets.

**Fig. 5.**
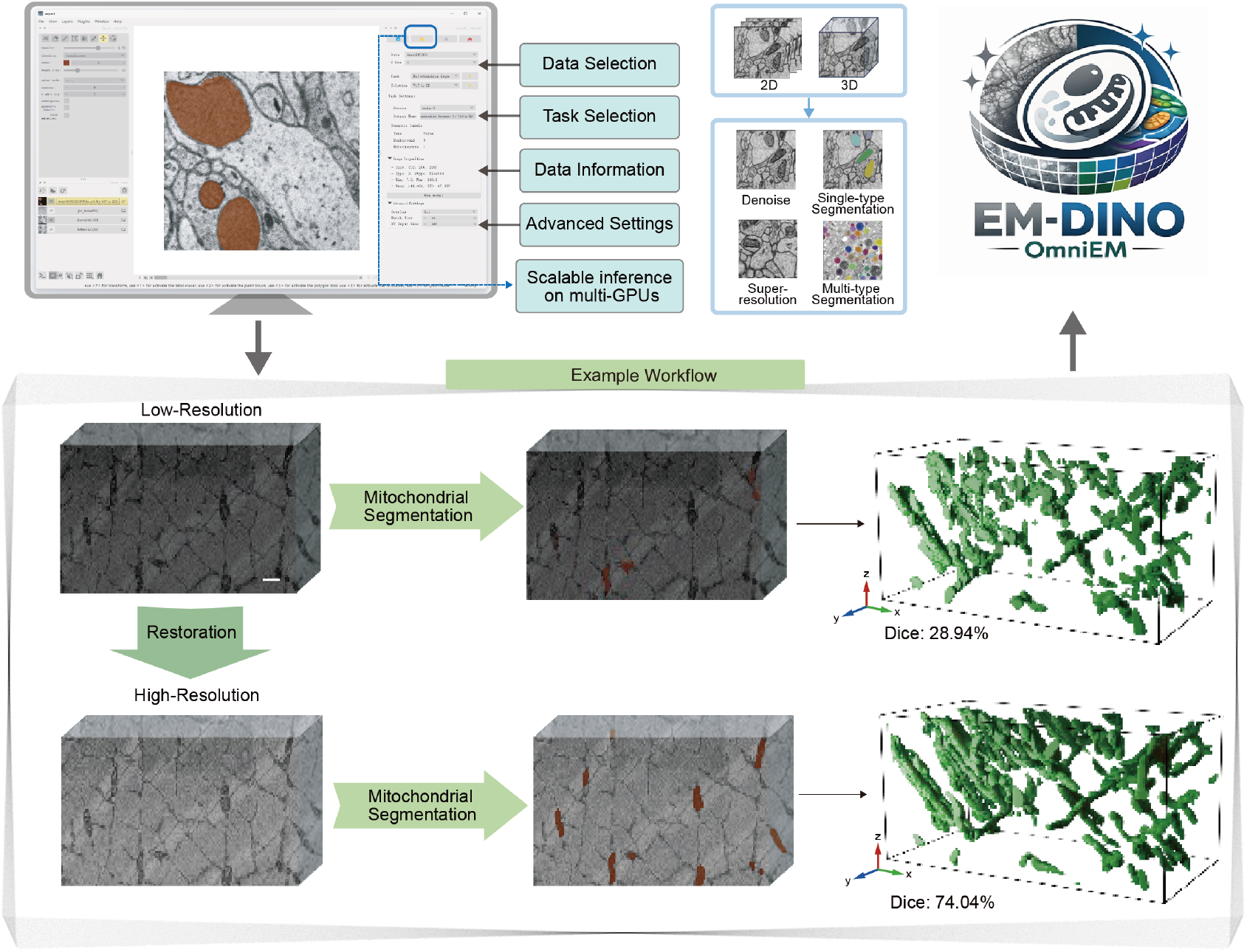
Integrated EM workflows enabled by OmniEM and the Napari Plugin. Screenshot of the OmniEM–Napari plugin interface (top right). The plugin supports interactive execution of multiple EM image analysis tasks (top middle). Middle and bottom: an example integrated workflow in which restoration is first applied to enhance image quality before segmentation, resulting in a substantial improvement in segmentation performance on the glycolytic muscle mitochondrial segmentation benchmark, from a segmentation-only Dice score of 28.94% (middle right) to 74.04% (bottom right). Scale bar, 1*µ*m.

## Results

### Evaluating Image Embeddings

To quantitatively assess the impact of EM-specific pretraining, we designed two complementary experiments. First, we used organ classification to test whether the learned embeddings can distinguish EM images from different organ types. Second, we examined how embeddings capture structure within a single EM image by dividing large-field images into smaller regions and visualizing their relationships. These experiments assess both differences across datasets and internal organization within individual EM images, supporting the ability of the pretrained embeddings to capture meaningful structural information about biological tissues and cells at the EM image level.

First, we evaluated image representations using two complementary organ classification tasks to assess how EM-specific pretraining influences embedding quality under different contextual conditions. The first, a context-aware task, used images from 11 organ types within *EM-5M* that retained rich morphological cues (Fig.2A). The second, a fine-grained task, employed small field-of-view images from *Histology Guide*^2^, which offer limited contextual information and present a more challenging generalization scenario for models (Fig.2D). In the context-aware task, all models exceeded 85% accuracy with just 10% of training data and surpassed 95% with full supervision, with all *EM-DINO* variants outperforming the *DINOv2* baseline (Fig.2B). Visualization of frozen embeddings further confirmed *EM-DINO’*s advantage, revealing well-separated, semantically coherent clusters compared to the more disorganized *DINOv2* embeddings (Fig.2C). In the more difficult fine-grained task, performance dropped across the board due to reduced context, yet *EM-DINO* maintained over 70% accuracy even with minimal supervision, outperforming all other models including *DINOv2*, which fell below 50%(Fig.2E). With increased training data, *EM-DINO* rapidly surpassed 90% accuracy and converged earlier than its counterparts. Consistent with this trend, two-dimensional UMAP visualizations of the embedding space showed clearer and more compact clusters for *EM-DINO* (Fig.2F), while t-SNE projections yielded similar results with improved class separation compared to other models (Extended Fig.3). Together, these results demonstrate that EM-specific self-supervised pretraining yields robust image embeddings that remain robust across a wide range of morphological and contextual variations in EM data.

Next, to assess whether image embeddings capture internal spatial organization within single EM images, we partitioned large-field EM images into overlapping patches, extracted embeddings, and projected them onto three principal components using PCA for RGB visualization (Extended Fig. 4A). In this representation, patches with similar embeddings appear with similar colors, enabling direct comparison between embedding organization and underlying ultrastructure shown in the corresponding large-field images and regions of interest. We applied this analysis to representative EM images from Histology Guide, including cardiac muscle^3^ and bone marrow megakaryocytes^4^. In cardiac muscle (Extended Fig. 4, B left), EM-DINO embeddings separated muscle from surrounding tissue and further resolved distinct myofibril sectioning patterns, corresponding to fibers cut in longitudinal versus cross-sectional orientations, consistent with known ultrastructural organization of sarcomeres and myofibrils in cardiomyocytes(Extended Fig. 4, C left),^33^. By contrast, DINOv2 embeddings failed to consistently distinguish muscle regions or capture these orientation-dependent differences (Extended Fig. 4D left). Similarly, EM-DINO delineated nuclear lobes in bone marrow megakaryocyte^34^, whereas DINOv2 produced fragmented and noisy representations(Extended Fig. 4, B D right). Together, these results demonstrate that EM-DINO embeddings preserve biologically meaningful spatial and contextual organization within large-field EM images across diverse tissue types.

Together, these experiments demonstrate that EM-DINO learns robust and transferable image representations, capturing both global differences across diverse EM datasets and fine-grained spatial organization within individual images. This capability positions EM-DINO as a general-purpose representation backbone for a wide range of downstream EM analyses.

### Robust Image Restoration with OmniEM

High-quality image restoration is fundamental to EM analysis, because high-throughput EM techniques often sacrifice resolution or dwell time (DT) for throughput, precluding reliable detection of fine ultrastructural features. Traditional restoration methods rely on regression, but regression-based models tend to oversmooth images and erase critical nanoscale details. More recently, a generative approach, EMDiffuse^15^, has shown promise; however, its diffusion-based process may introduce hallucinated artifacts that mislead downstream analyses. To overcome these challenges, we implemented both denoising and super-resolution within the OmniEM framework, incorporating embeddings extracted from a pretrained EM-DINO model. We evaluated its performance by comparing it with EMDiffuse^15^ and PSSR^35,36^, representing diffusion-based and regression-based approaches, respectively. From the results, OmniEM achieves comparable or superior performance to PSSR and EMDiffuse in denoising and super-resolution tasks (Fig. 3).

Specifically, we introduce a loss function based on embeddings derived from EM-DINO representations to encourage the preservation of fine structural details while mitigating over-smoothing. This design enables OmniEM, a primarily regression-based network, to recover high-resolution features with greater fidelity than existing methods. Notably, EM-DINO’s embedding-based loss contributes to the observed performance gains of OmniEM (Extended Data Fig. 5). We introduce two hyperparameters, *α* for image-level embedding loss and *β* for patch-level embedding loss, to regulate the strength of this supervision during restoration training. Incorporating the embedding loss markedly reduces unwanted smoothness and enhances image details in the denoised output, especially at high noise levels (Extended Data Fig. 5A). For example, under high-noise conditions, the Learned Perceptual Image Patch Similarity (LPIPS) on denoised images drops from 0.575 (without embedding loss) to 0.211 (with *α* = 0.25, *β* = 0), to 0.305 (with *α* = 0, *β* = 0.25), and to 0.204 (with *α* = 0.25, *β* = 0.25). Lower LPIPS values indicating closer agreement with the ground truth in terms of human-perceived image quality. This improvement reflects the contribution of image-level embedding and patch-level embedding to enhanced perceptual fidelity. Further increases *α* and *β* beyond 0.25 do not significantly change LPIPS or subjective perceptual quality any more. For consistency, we applied the same hyperparameter values (*α* = 0.25 and *β* = 0.25) to all the following restoration experiments.

In the denoising task, the three models exhibit distinct restoration behaviors (Fig. 3), with differences becoming particularly pronounced under high-noise conditions (Fig. 3C): PSSR preserves general morphology but misses structural details; EMDiffuse recovers visually plausible textures but include spurious artifacts not found in the long-DT ground truth; in contrast, OmniEM conservatively restores key cellular topologies with minimal spurious structural artifacts, preserving structural consistency essential for downstream tasks. The artifacts during restoration can compromise the validity of downstream analyses. For example in a vesicle segmentation task(Extended Data Fig. 6 A), a nnU-Net tends to detect more vesicles from EMDiffuse images, but with a high false-positive rate. In contrast, although OmniEM yields fewer detections (reflecting a more conservative restoration), the majority of its detected instances are accurate, resulting in substantially higher precision. These observations are corroborated by precision–recall and ROC analyses (Extended Data Fig. 6 B and C), in which OmniEM achieves larger areas under the curves, indicating more accurate and reliable segmentation.

To quantitatively assess restoration quality, we evaluated perceptual similarity (LPIPS), structural similarity (Feature Similarity Index Measure, FSIM), and image resolution both using ground-truth–independent (decorrelation-based) metrics and ground-truth–dependent (Fourier ring correlation, FRC), consistent with metrics used in EMDiffuse. (Fig. 3B and D). Under all noise levels, OmniEM achieved the lowest LPIPS values among the three models (OmniEM < EMDiffuse < PSSR), indicating the highest perceptual fidelity to the ground truth (Fig. 3B and D and Extended Data Fig. 7). For FSIM, OmniEM exhibited best scores under high- and medium-noise conditions (EMDiffuse < PSSR < OmniEM)(Fig. 3 D and Extended Data Fig. 7), reflecting its ability to preserve structural details without excessive smoothing. However, under low-noise conditions(Fig. 3 B), PSSR achieved better results,possibly because its regression-based formulation is sufficient when the input images already contain rich fine structural details. Decorrelation-based resolution (Fig. 3B and D and Extended Data Fig. 7), which assesses structural sharpness independently of ground truth, showed intermediate values for OmniEM (EMDiffuse < OmniEM < PSSR), consistent with the diffusion process facilitating the generation of high-frequency structural information. However, FRC-based analysis showed that OmniEM surpassed EMDiffuse (Fig. 3B and D and Extended Data Fig. 7), suggesting that OmniEM could recover more ground-truth-related details. Unexpectedly, PSSR achieved the highest FRC-based resolution, likely because the regression-based design smooths the entire image, making it less affected by noise. A similar phenomenon was observed when *α* and *β* were set to 0, yielding higher FRC-based resolution than that obtained with *α* and *β* were set to 0.25 (data not shown). Combining the two resolution metrics, OmniEM exhibited intermediate numerical resolution values, while qualitatively achieving a balance between the over-generation of EMDiffuse and the over-smoothing of PSSR. Paired comparisons across image tiles (lines between scatters) and statistical analyses confirmed that OmniEM provided a robust compromise between perceptual accuracy and structural preservation.

Beyond denoising, OmniEM exhibits consistent performance in super-resolution (Fig. 3E–H), comparable to its behavior in the denoising task. OmniEM reconstructs critical subcellular details from low-resolution inputs under high-noise (Fig. 3E), low-noise (Fig. 3G), and medium-noise (Extended Data Fig. 7C). By contrast, PSSR yields overly smoothed results while EMDiffuse recovers fine details but occasionally introduces artificial textures absent from the high-resolution reference. For example, in Fig. 3G, EMDiffuse produces a membrane-like structure extending from the central mitochondrion that, while visually consistent with biological texture, lacks physiological meaning and is absent in the reference image. By comparison, OmniEM enhances ultrastructural features relative to PSSR while avoiding the spurious artifacts observed in EMDiffuse. Quantitative evaluation across image tiles consistently supports the trends observed in the qualitative analysis. (Fig. 3F and H). OmniEM consistently outperforms EMDiffuse in LPIPS, FSIM and FRC-based spatial resolution metrics, which emphasize perceptual, feature-related, and structural accuracy relative to reference images. It also significantly surpasses PSSR in decorrelation-based resolution, which reflects the intrinsic spatial resolution of the images themselves. Paired comparisons and statistical analyses further demonstrate that OmniEM consistently balances detail enhancement with faithful reconstruction, yielding super-resolved images that are both structurally accurate and perceptually closer to the high-resolution reference.

In conclusion, OmniEM delivers image restoration performance comparable to previous well-performing methods while avoiding model over-interpretation that could mislead downstream analyses, especially under high-noise. By producing more conservative and biologically plausible reconstructions, OmniEM provides the reliability needed for downstream tasks that depend on accurate ultrastructural representation. The restoration performance could be further improved by exploring enhanced architectural designs or loss functions.

### Precise image segmentation

As expert annotations are costly and limited to specific datasets, accurate organelle segmentation forms the basis for quantitative ultrastructural analysis and biological discovery, while existing methods struggle to generalize across diverse EM datasets and semantic categories. To explicitly evaluate this capability, we employed mitochondrial segmentation tasks and a multi-class segmentation task, with training and testing data drawn from different EM volumes to test generalization to unseen data. Further, we benchmark OmniEM on 3D EM segmentation to evaluate how 2D-pretrained EM-DINO embeddings transfer to volumetric settings. Since EM-DINO is trained only on 2D images, a naïve baseline applies slice-wise inference by cropping volumes along the z-axis and mapping 2D predictions back into 3D space. Beyond this baseline, OmniEM supports volumetric training by integrating limited z-axis context within its intermediate feature blocks while still relying on 2D-pretrained embeddings. Accordingly, we report two settings: 2D experiments, with 2D training/inference and 3D volume evaluation, and 3D experiments, with both training and inference performed on 3D volumes. Again, these findings highlight the model’s potential as a general-purpose segmentation framework for EM images.

Mitochondrial morphology and distribution serve as critical indicators of cellular metabolic state and disease progression. However, the substantial morphological heterogeneity of mitochondria across tissues poses a major challenge for cross-dataset generalization in automated segmentation models^37^. In this work, we evaluated OmniEM on two complementary benchmarks. The first is the CEM-MitoLab dataset, which comprises labeled 2D EM images from multiple sources and is used to assess 2D training with evaluation on seven external 2D and 3D datasets withheld from training ^13^. The second benchmark is MitoEM^38^, which provides two annotated EM volumes from different species (rat and human). We trained models on rat sub-volumes and evaluated on the human volume, forming a cross-species challenging 3D training and 3D evaluation setting. For comparison, we include MitoNet^13^ as a 2D-only method and MedNeXt^39^ as a 3D-only model, while nnU-Net^40^ and OmniEM are evaluated under both 2D and 3D training/inference conditions.

In results, the publicly released MitoNet checkpoint maintains strong segmentation accuracy across all seven benchmarks on the CEM-MitoLab, as expected(Figure 4A, third row; light-apricot histogram bars in Figure 4B). OmniEM, however, achieves comparable or superior performance to MitoNet on five of seven datasets (Figure 4A, below row; light-blue histogram bars in Figure 4B)), highlighting the value of EM-DINO pretraining in providing robust, transferable embeddings for mitochondria segmentation. Overall, OmniEM gives the best average dice 78.77%, compared to MitoNet (71.87%) and nnU-Net (63.34%) (Figure 4B). The largest improvement occurs on the salivary gland dataset, where mitochondria exhibit flat, bowl-shaped geometries, high organelle density, and weak contrast against salivary granules (Figure 4A, sixth column). In this particularly challenging case, OmniEM’s Dice score leaps from 11.27 % (MitoNet) to 42.59 % (Figure 4B, sixth inset), demonstrating that EM-DINO pretraining confers enhanced representational capacity for segmenting rare or low-contrast mitochondrial morphologies. On the 3D MitoEM benchmark, OmniEM achieved the highest performance (Dice = 93.63 %), outperforming nnU-Net (Dice = 90.23 %) and MedNeXt (Dice = 87.82 %) (Extended Figure 9A & C). These results underscore OmniEM’s strong cross-dataset generalization performance, even on unseen tissue types and diverse imaging conditions.

Compared to single-organelle segmentation, multi-organelle segmentation is considerably more challenging due to the diverse morphologies, sizes, and close spatial relationships among different organelles. Variability in imaging conditions, staining quality, and the inherent ultrastructural complexity further complicate accurate delineation, often resulting in overlapping boundaries and ambiguous separations between adjacent structures. To address these challenges, we implemented a multi-class segmentation head within OmniEM and evaluated its performance on the BetaSeg dataset^19^, which includes annotations for Golgi apparatus, mitochondria, insulin secretory granules, and nuclei (all other pixels are treated as background). We further evaluated OmniEM under a zero-shot generalization setting by splitting the BetaSeg volumes into disjoint training (high_c1, high_c2, high_c3) and testing (high_c4, low_c1, low_c2, low_c3) sets. To assess robustness across dimensional settings, we conducted both 2D and 3D experiments by training on cropped 2D slices or 3D subvolumes, respectively. For baseline comparison, we selected GobletNet^41^, a recently 2D-only model that uses wavelet transforms to extract high-frequency features to improve segmentation accuracy, MedNeXt for 3D segmentation^39^, and nnU-Net^40^ for both 2D and 3D settings. In addition, for 2D experiments, we subsampled the training data into four fractions, 1 %, 30 %, 60 %, and 100 %, to test few-shot learning capability.

In results, OmniEM consistently outperforms GobletNet across all 2D training fractions, with nnU-Net providing a strong baseline(4). Notably, under the extreme few-shot setting (1 % training data), both OmniEM and nnU-Net generalize well to unseen volumes, achieving mean Dice scores above 70 %, whereas GobletNet remains substantially lower. OmniEM further attains the highest overall performance in this regime, particularly improving segmentation of Golgi apparatus, mitochondria, and nuclei. Across increasing data availability (30–100 %), OmniEM maintains stable and consistent gains over nnU-Net in mean Dice, with pronounced improvements for nuclei and mitochondria. Such a gap between GobletNet and the other models likely reflects our deliberate zero-shot training strategy, in which models are trained only on “high” series volumes and evaluated on completely separate “low” series volumes that differ in imaging characteristics. On the 3D BetaSeg benchmark, OmniEM (mean Dice = 83.96%), nnU-Net (84.63%), and MedNeXt (83.72%) show comparable performance across all organelle classes (Source table 3 and Extended Data Fig. 9B & D). Together, these results demonstrate that OmniEM, leveraging EM-DINO embeddings, not only improves generalization and data efficiency in 2D tasks but also provides a robust and flexible framework for volumetric EM segmentation.

These results highlight OmniEM’s strong generalization and data efficiency, enabled by the rich representational capacity inherited from EM-DINO pretraining. Without relying on handcrafted features or explicit task-specific priors, OmniEM effectively captures diverse organelle morphologies and subtle ultrastructural cues. This underscores the potential of EM foundation models to deliver robust, versatile segmentation across complex electron microscopy datasets.

### Integrating OmniEM into Scalable EM Workflows

OmniEM’s multi-task design enables streamlined, end-to-end EM workflows by combining advanced restoration, super-resolution, and segmentation within a single, user-friendly framework. We have packaged OmniEM’s restoration and segmentation modules into an interactive Napari plugin (Fig. 5), allowing users to apply OmniEM via a graphical interface. To accommodate the diverse characteristics of EM datasets, including small- or large-field images and both 2D and 3D modalities, we provide two inference modes in the OmniEM–Napari plugin. The first supports in-memory, single-GPU inference for small-scale datasets, whereas the second enables local, multi-GPU inference for large-scale slices and volumes by leveraging Dask-based lazy loading instead of fully loading data into memory. These complementary modes allow users to flexibly load data, adjust parameters according to available hardware resources, and interactively visualize results through the Napari graphical interface. Using the glycolytic muscle mitochondrial segmentation benchmark as an example, we contrast this integrated workflow with the conventional paradigm that focuses on isolated processing steps, where restoration and segmentation are performed using separate models. In contrast, the OmniEM plugin supports a more realistic end-to-end workflow by sequentially combining restoration and segmentation on the same dataset, resulting in a substantial improvement in segmentation performance in this dataset, with the Dice score increasing from 28.94% to 74.04% when restoration is applied prior to segmentation. Similar workflows can be constructed for other datasets and tasks using the same plugin interface. By bundling pretrained EM-DINO and OmniEM checkpoints within this plugin, we provide a comprehensive and reproducible workflow for scalable, high-quality EM analysis with minimal technical overhead.

## Discussion

In this work, we demonstrate that large-scale, EM-specific self-supervised learning enables the construction of a foundational vision model capable of unifying previously fragmented EM workflows. At the core of our approach is a novel data-curation pipeline that systematically deduplicates and filters vEM patches, reducing redundancy while keeping biological and technical diversity. By pairing EM-DINO for robust feature extraction with OmniEM for flexible downstream prediction tasks, we deliver a suite of tools tailored to the EM community’s demand for efficient, scalable, and reproducible analysis. Although achieving maximizing benchmark scores was not our primary objective, OmniEM nonetheless attains strong performance in denoising, super-resolution, and segmentation without reliance on handcrafted heuristics or specialized architectures, highlighting the generalization power of EM-DINO’s learned representations.

### Methodology

A central but often underappreciated component of this work is the data curation strategy underlying EM-5M. Volume electron microscopy data are characterized by extreme redundancy, with neighboring slices often differing only marginally at the pixel level. To address this, we adopted an iterative data reduction recipe: an initial EM-DINO model was pretrained and used to identify and suppress redundant image content, after which newly added data were incorporated and the model retrained. As a result, EM-5M should not be interpreted as merely a collection of 5 million images, but rather as a distilled representation derived from more than 500 TB of raw EM data, retaining high structural diversity while mitigating redundancy. Data diversity was treated as a primary design constraint. Although available EM data are heavily skewed toward brain tissue (driven by connectomics) and mouse samples (notably from OpenOrganelle), we explicitly constrained any single domain to contribute no more than 50% of the dataset and excluded several large but highly homogeneous sources (e.g., MICrONS^20^) to avoid overdominance. Importantly, we empirically evaluated pretraining on substantially larger but unreduced datasets and observed inferior downstream performance, indicating that scale alone is insufficient; rather, controlling redundancy and promoting structural diversity are critical factors for effective EM representation learning. Together, these observations suggest that data recipe (how EM data are selected, balanced, and distilled) are as important as model architecture or training objective for building generalizable EM foundation models.

Image representation learning, often realized through large-scale self-supervised pretraining, aims to extract meaningful and structured information to support multiple downstream analyses^42^. In natural image analysis, large-scale self-supervised learning frameworks such as MoCo^43^ and SimCLR^44^ have demonstrated that a single pretrained representation can generalize effectively across a wide range of downstream tasks, bridging traditional divides between recognition, segmentation, and reconstruction. These approaches highlight the value of learning transferable, task-agnostic visual embeddings from diverse data. In contrast, progress in the EM domain has largely followed task-specific trajectories. Most prior self-supervised methods, including CEM^13,45^, SegNeuron^46^, and MASTER^47^, were designed to enhance performance for particular segmentation benchmarks, rather than to establish a unified representation capable of supporting multiple EM analysis tasks. In this work, we build upon DINOv2 as the pretraining strategy, as it combines masked image modeling with contrastive self-distillation and has demonstrated strong transferability across vision tasks.

The EM-DINO/OmniEM framework employs a hybrid pretraining–training strategy: EM-DINO performs purely 2D self-supervised pretraining to learn generalizable image representations, while OmniEM uses a hybrid architecture that combines 2D encoders with task-specific 2D or 3D decoders, enabling both 2D and 3D downstream applications. In contrast, most existing self-supervised frameworks operate in a single data dimensionality: many are strictly 2D, as in natural images^43,48^, whole-slide imaging^49^, and EM-specific methods^13,47^, whereas others are fully 3D for biomedical image pretraining^46,50^. Our mixed design reflects the nature of EM data, which spans both slice-based 2D analyses and volumetric 3D tasks, yet often lacks well-curated and fully aligned 3D representations. Many EM workflows remain inherently 2D, including image alignment^51^ and subcellular anatomy studies^52^, while volumetric data are common in fields requiring large-scale analysis, such as connectomics^6^. Accordingly, EM-DINO/OmniEM operates directly on raw 2D EM images without needing complete 3D volumes, yet effectively transfers to 3D tasks through OmniEM. This design positions EM-DINO/OmniEM as a flexible and generalizable representation learning framework that complements existing 2D- or 3D-specific approaches and accommodates the heterogeneous data formats typical of EM studies.

### Downstream tasks

EM restoration plays a pivotal role in EM-based research and is a critical step toward enabling high-throughput EM imaging. Recent advances in deep learning–based methods have substantially outperformed conventional image restoration approaches^53^, but challenges remain. A key issue is balancing high-quality restoration with image fidelity. Regression-based models, such as PSSR^35,36^, are effective at suppressing noise but often lead to over-smoothing, resulting in the loss of fine ultrastructural details. Conversely, diffusion-based methods pioneered by EMDiffuse^15^ address over-smoothing but tend to over-interpret the data, introducing visually plausible yet physiologically incorrect structures. In the absence of reference images, such artifacts are difficult to distinguish and may lead to erroneous biological conclusions. In this work, we address this by incorporating embeddings from pretrained EM-DINO into the training loss, achieving a practical balance between detail recovery and reconstruction fidelity across various noise levels. This strategy enables OmniEM to restore fine ultrastructural features while avoiding spurious structural artifacts, improving the reliability and biological plausibility of the reconstructed images. In parallel, our restoration results also highlight limitations in evaluation metrics for EM. Commonly used pixel-based metrics for natural images, such as PSNR and SSIM, have been shown to be inadequate for assessing EM image quality^15^. Although perceptual and structural metrics such as LPIPS and FSIM provide more informative assessments, they are not specifically designed for EM data. For example, LPIPS relies on feature representations learned from natural images, and the substantial domain gap between natural and EM images limits the appropriateness of directly applying such models. We believe EM-DINO can provide a promising starting point for EM-specific evaluation. In short, developing robust metrics and perceptual models tailored to EM remains an important direction for future work.

EM segmentation has traditionally been dominated by U-shaped convolutional architectures and their variants, with recent work exploring attention- and state-space–based extensions that primarily target improved performance under matched training and testing conditions^14,54,55^. In contrast, this study places particular emphasis on robustness under distribution shift, which is a pervasive challenge in practical EM analysis. On the BetaSeg benchmark, where training and testing data originate from the same imaging series and species, OmniEM performs comparably to established pipelines such as nnU-Net. However, on cross-dataset and cross-species mitochondrial benchmarks spanning multiple datasets, imaging conditions, and species, OmniEM exhibits more stable performance. We attribute this behavior not to architectural novelty alone, but to the use of EM-specific self-supervised pretraining combined with a lightweight task-adaptation architecture, which encourages the model to learn transferable ultrastructural features rather than task- or dataset-specific cues. This distinction is especially important in EM, where expert annotations are expensive and limited, and models are frequently applied to unlabeled data that differ substantially from the available training distributions. By integrating self-supervised representation learning with supervised task adaptation, the EM-DINO/OmniEM framework offers a practical balance between in-distribution performance and cross-dataset generalization, addressing a central bottleneck in scalable EM image analysis.

### Limitations and future extensions

Because most modern image processing and interpretation pipelines hinge on the availability of suitable features, EM-DINO can serve as a universal feature extractor, readily integrated into existing tools to enhance performance. For example, in image stitching and alignment, point correspondence detection often depends on classical descriptors such as SIFT^56^, or more recently, deep encoders trained to suppress artifacts in EM images^16,20^. EM-DINO’s semantically enriched features could significantly improve matching accuracy across sections, particularly under varying imaging conditions. Similarly, in connectomics pipelines that require the linking of neurite fragments, cellular identity assignment, and synapse detection, practitioners typically rely on separate classifiers optimized for each task^57^. We propose that EM-DINO can serve as a unified representation backbone across these components, reducing task fragmentation and simplifying pipeline integration.

While EM-DINO provides versatile and semantically rich features for diverse tasks, representation learning alone does not equate to full biological understanding of EM data. EM images span large fields of view and volumetric contexts, and image-only self-supervised pretraining lacks explicit biological semantics, such as organelle or cellular identities^2^. We therefore view representation learning as a foundational step toward large-field EM image understanding and semantic alignment across datasets and modalities. By providing open models and tools for learning EM representations, this work establishes a basis for future integration with higher-level biological inference. For example, strategies for aggregating local 2D image embeddings could be used to construct global representations of large-field images, similar to approaches previously applied in whole-slide imaging^58^. Beyond EM alone, EM-DINO could serve as a modular component for multimodal integration, linking EM data with complementary modalities such as cryo-electron tomography, light microscopy^59^ or even natural language annotations. Taken together, these directions highlight the potential of EM-DINO as a foundational image-only feature extractor for EM data, paving the way toward more comprehensive and integrative analyses of cellular ultrastructure.

Looking ahead, several strategies could further strengthen EM-DINO’s utility, improving both its generalization across diverse EM datasets and its computational efficiency. First, expanding EM-5M beyond its current emphasis on brain and mouse vEM to include underrepresented organs (e.g., kidney, heart, bone) and non-mammalian species will ensure downstream models learn truly transferable representations across diverse EM applications. To facilitate this, we have made our data curation tools publicly available, enabling the community to contribute additional datasets and further enrich EM-5M. Second, although EM-DINO uses a Vision Transformer backbone, alternative architectures, such as hierarchical transformers like Swin Transformer^60^ or Mamba^61^, may better capture EM’s anisotropic resolution and high noise levels. Systematic benchmarking of these backbones on EM-specific tasks will clarify which design choices maximize ultrastructural detail. Finally, EM-DINO’s large-scale pretraining yields robust embeddings, but fine-tuning remains computationally intense; parameter-efficient techniques like Low-Rank Adaptation (LoRA)^62^ offer a promising route to rapid adaptation on new datasets with minimal overhead. Additionally, OmniEM currently employs separate branches for restoration and segmentation, missing potential synergies from shared features. A multi-task or auxiliary-learning framework could unify these tasks, improving both efficiency and accuracy^63^, and moving closer to a truly general-purpose, dense-prediction model for EM data. By addressing these challenges, future work can build on our EM-specific foundation to create ever more versatile, accurate, and efficient tools for nanoscale biological discovery.

## Data and code availability

We maintain the project repository on GitHub, providing public access along with comprehensive documentation for users^5^.

## Acknowledgements

We thank Quanyue Ma for assistance in downloading EM images from HistologyGuide.

## Author contributions statement

These authors contributed equally: Liuyuan He, Ruohua Shi, and Wenyao Wang.

## Contributions

L.M. and W.W. supervised the research. L.M., W.W. and L.H. conceived of the technique. W.W., L.H., R.S., G.F. and Y.C. implemented the algorithm and conducted the experiments. L.H. curated the pretraining dataset, pretrained backbone models, designed the framework of OmniEM, and developed OmniEM-Napari. R.S. implemented the downstream tasks and organized the codes and models. All authors had access to the study and wrote the paper.

## Extended data

Source Data 1: Performance comparison of Organ Classification Tasks (Accuracy in %) Source Data 2: values of metrics in figure 3

Source Data 3: values of metrics in segmentation tasks, represented in figure 4 and Extended Data figure 9

Source Data 4: values of metrics in Extended figure 5

Source Data 5: values of metrics in Extended figure 6

Source Data 6: values of metrics in Extended figure 7 Extended Data Table 1: Sources of datasets

**Extended Data Fig. 1.**
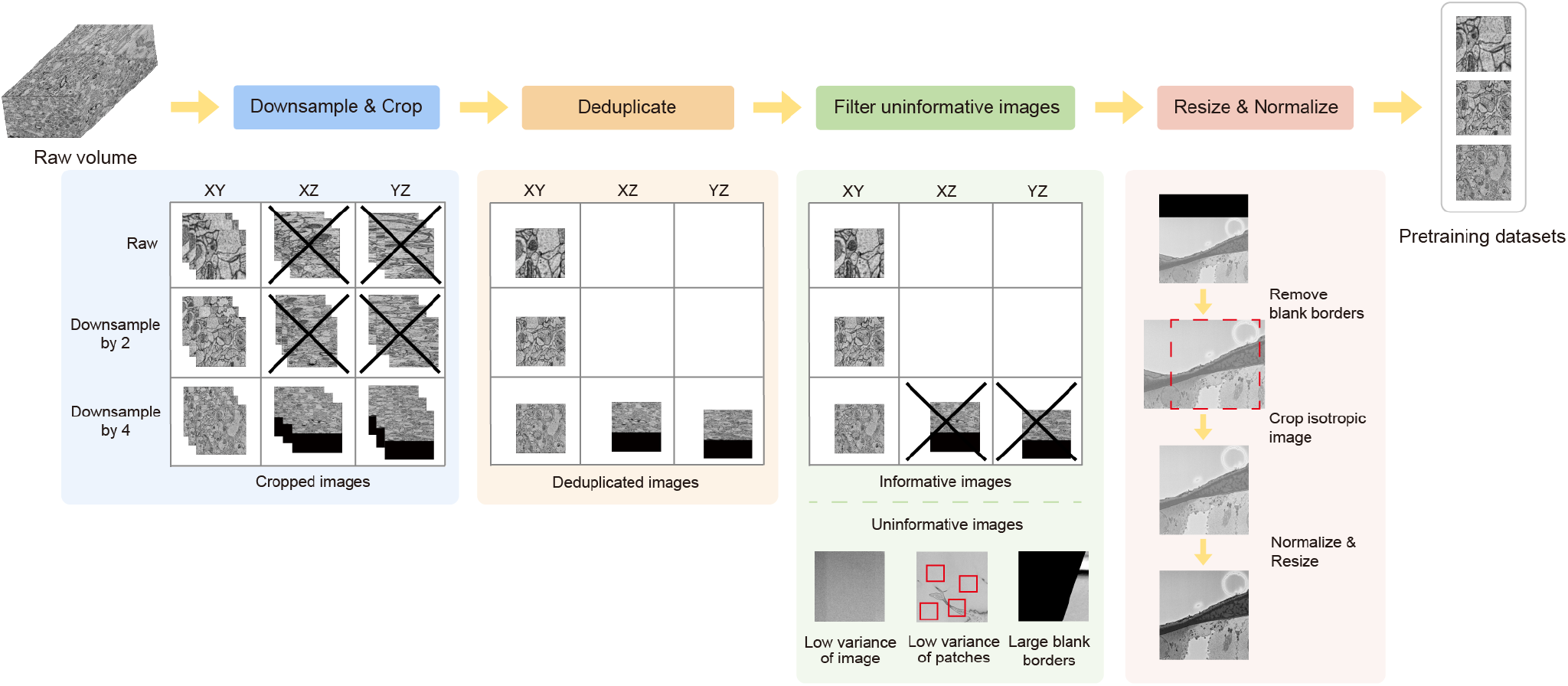
EM-5M data curation pipeline. Volumetric datasets are processed through the following steps: standardization, downsampling, cropping, deduplication, uninformative region filtering, and patch-level resizing and normalization. For 2D datasets, all steps are applied except for deduplication.

**Extended Data Fig. 2.**
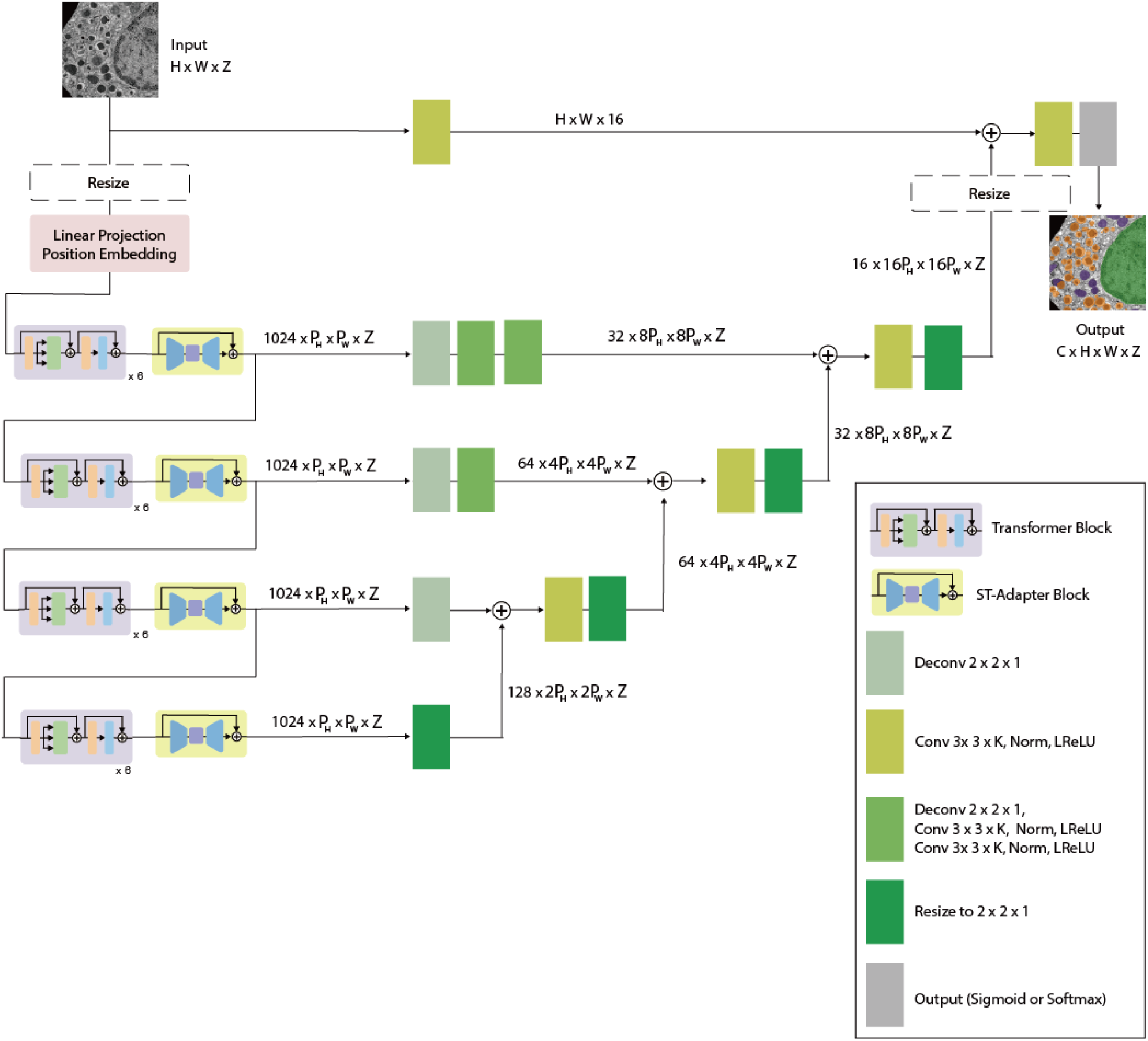
OmniEM architecture. OmniEM adopts a U-shaped encoder–decoder architecture with task- and dataset-dependent variations indicated by internal variables. Two alternative resize locations are shown (before linear projection or after the decoder), although only one is used per configuration. The spatial feature sizes *P*_*H*_ and *P*_*W*_ depend on the chosen resize strategy. All convolutional layers use 3 *×* 3 *×K* kernels, with *K* = 1 for 2D and *K* = 3 for 3D inputs.

**Extended Data Fig. 3.**
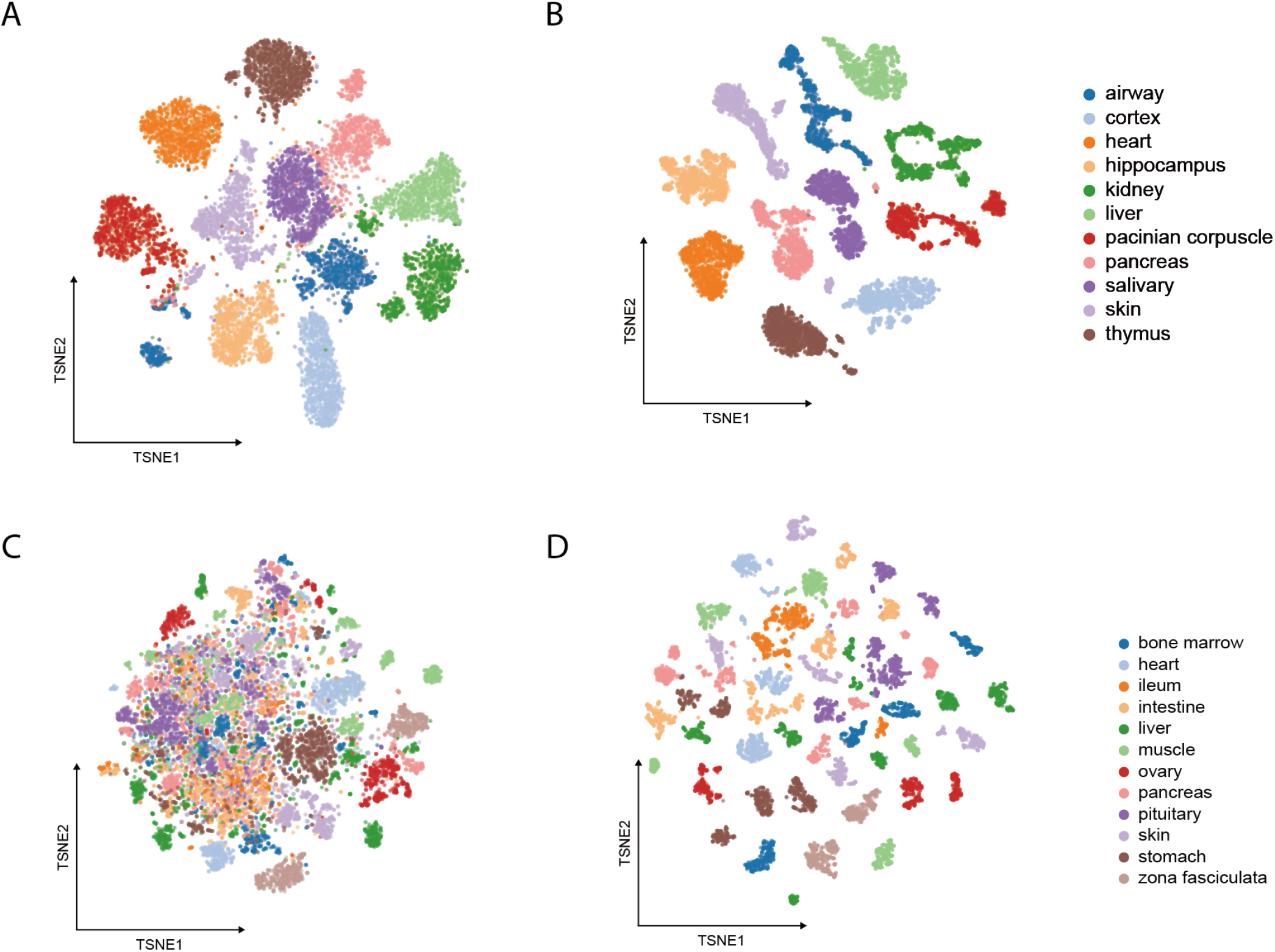
The t-SNE visualization of embedding spaces for organ classification. (A) Image embeddings from the context-aware classification task, corresponding to Fig. 2C. (B) Image embeddings from the fine-grained classification task, corresponding to Fig. 2F.

**Extended Data Fig. 4.**
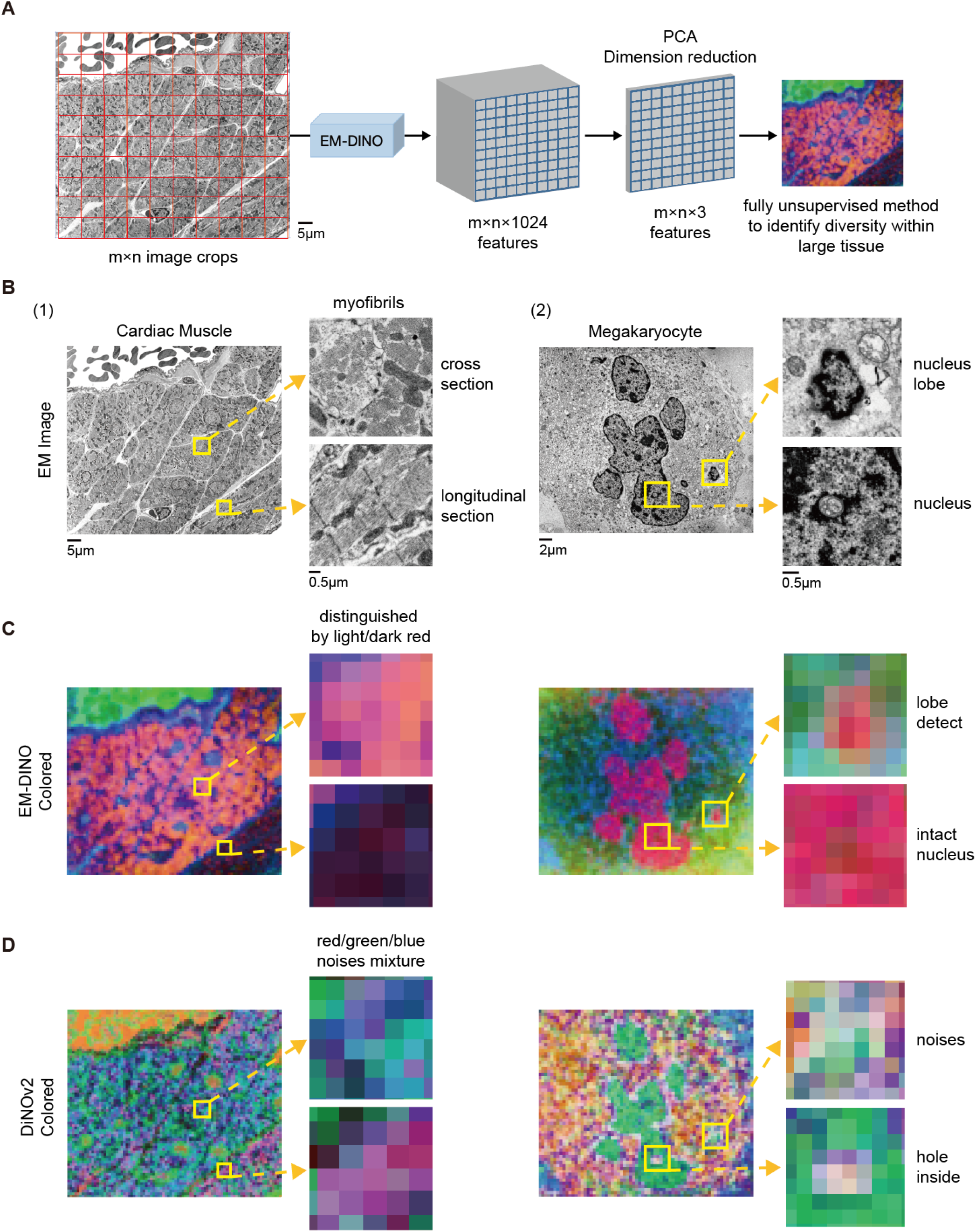
Spatial organization of image embeddings. **(A)** Schematic illustrating the visualization pipeline for large-field EM images, in which overlapping sub-regions are cropped, embedded, and projected for spatial analysis. **(1) Cardiac muscle**. Large-field EM image of cardiac muscle, with regions of interest (ROIs) highlighting myofibrils sectioned in cross-sectional and longitudinal orientations. **(2) Bone marrow megakaryocyte**. Large-field EM image of a megakaryocyte, with ROIs showing lobulated nuclei surrounded by cytoplasm and an intact nucleus. **(B)** Large-field EM images and corresponding zoomed-in ROIs. **(C)** PCA-based RGB visualization of EM-DINO embeddings extracted from overlapping patches. EM-DINO separates distinct myofibril sectioning patterns in cardiac muscle, resolves nuclear lobes, and distinguishes intact nuclear regions in megakaryocytes. **(D)** Corresponding PCA visualization using DINOv2 embeddings. DINOv2 fails to resolve myofibril orientation and nuclear lobulation, producing mixed and noisy color patterns with discontinuities within nuclear regions. Scale bars range from 0.5*µ*m to 5*µ*m.

**Extended Data Fig. 5.**
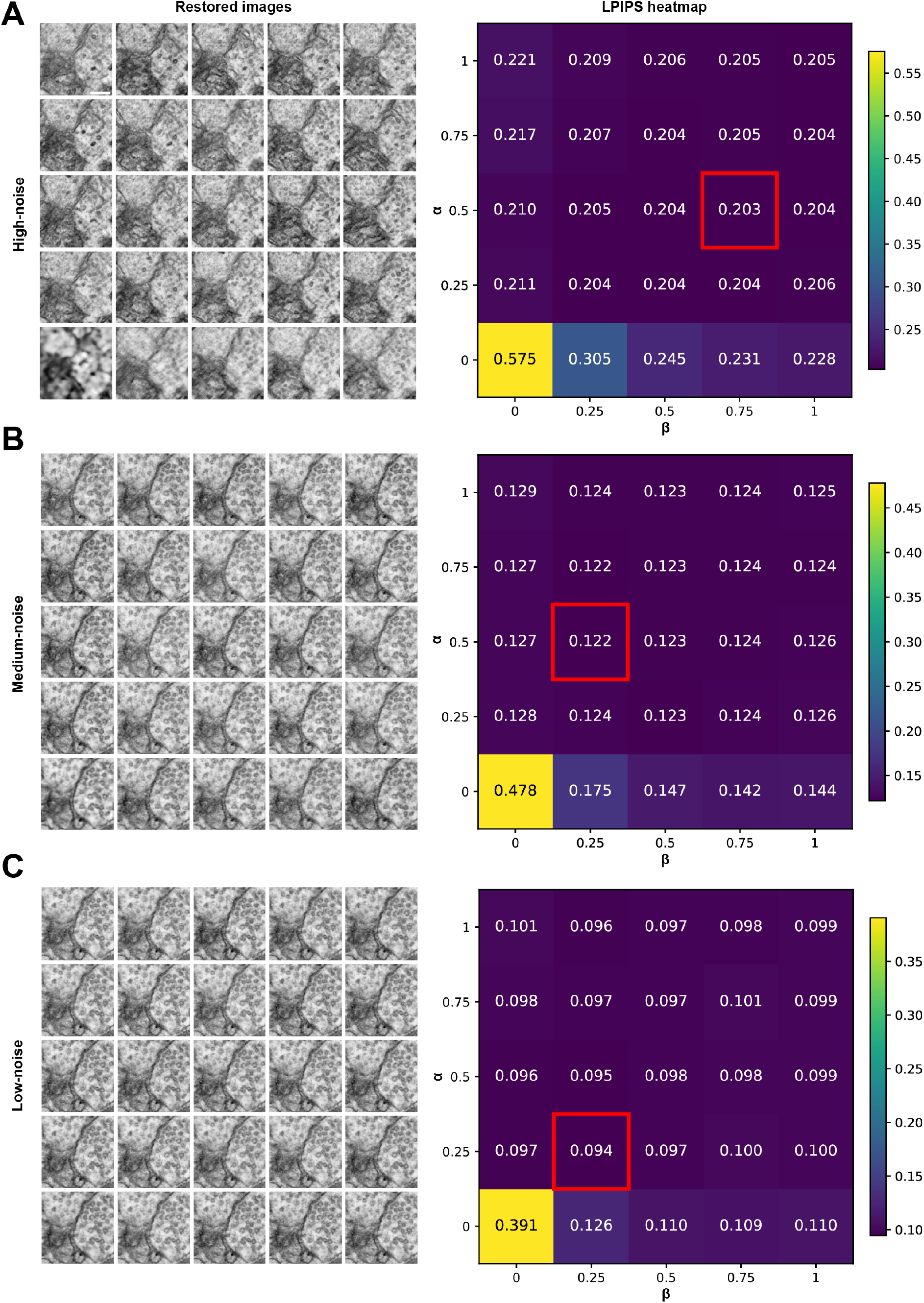
Grid search of image-level and feature-level embedding weights for denoising performance. **(A-C)** Illustration of the effects of image-level and feature-level embedding weights on denoising performance under high-, medium-, and low-noise conditions, respectively. For each noise level, the left column shows restored images from the same input image, and the right column shows the corresponding LPIPS heatmap. In the image grid, the bottom-left corner represents the case in which both embedding weights are set to zero. The image-level embedding weight (*α*) increases from 0 to 1 along the vertical axis, whereas the feature-level embedding weight (*β*) increases from 0 to 1 along the horizontal axis. LPIPS heatmaps follow the same axis orientation. The best LPIPS value in each heatmap is outlined with a red box. Scale bar: 200 nm.

**Extended Data Fig. 6.**
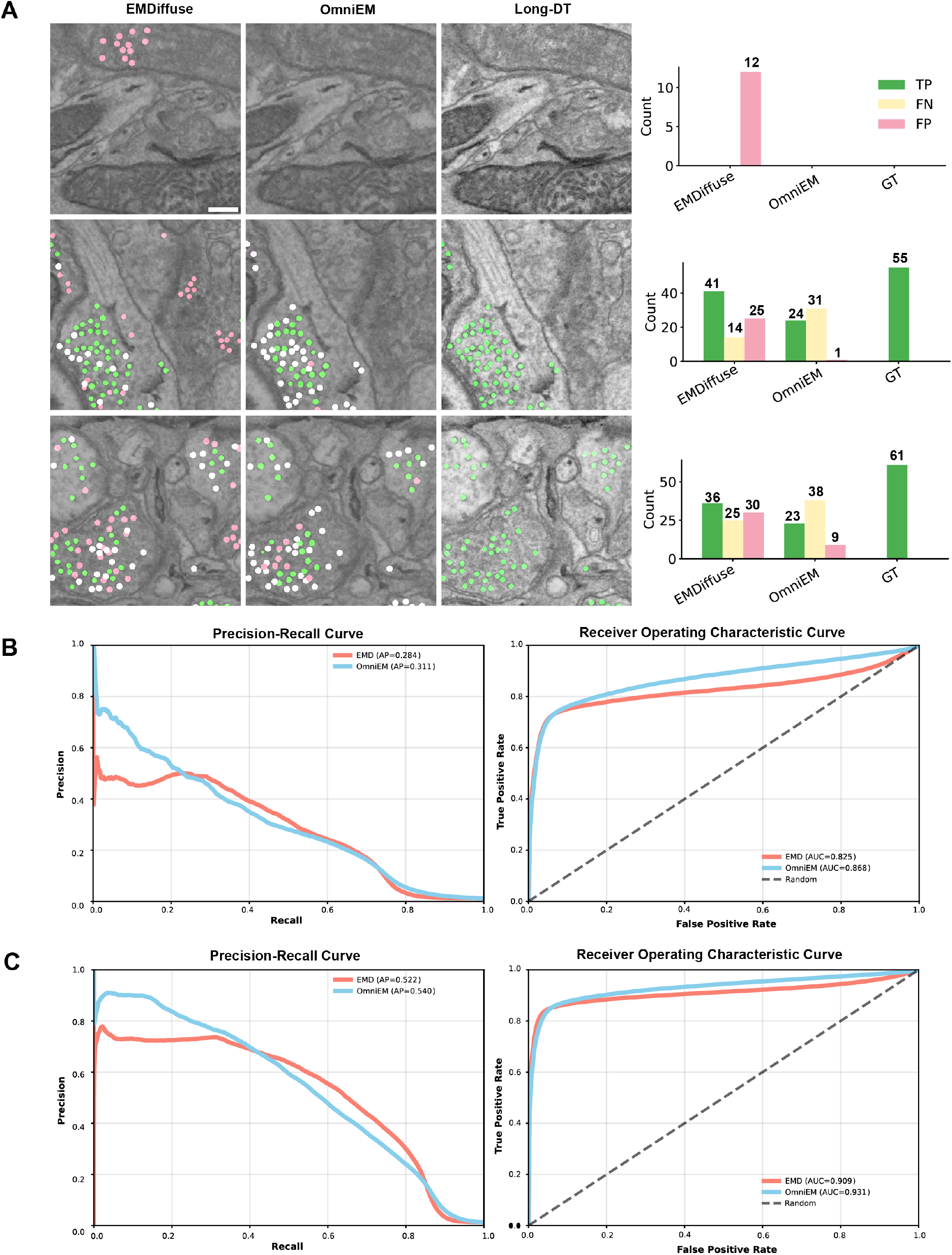
Comparison of vesicle detection performance between EMDiffuse and OmniEM restoration images. **(A)** Vesicle segmentation results produced by nnU-Net on images restored by EMDiffuse and OmniEM from high-noise input, as well as on the reference (Long-DT) images. Vesicles that are consistent with the Long-DT images are shown in green (true positives,TP), vesicles present in the Long-DT images but missed in the model restored images are shown in yellow (false negatives, FN), and vesicles absent in the Long-DT images but incorrectly detected are shown in pink (false positives, FP). The corresponding vesicle counts are displayed on the right. **(B)** Precision–recall (PR) and receiver operating characteristic (ROC) curves comparing OmniEM and EMDiffuse under high-noise conditions (one image montage; xy dimensions, 1596 × 1596 pixels). **(C)** Precision–recall (PR) and receiver operating characteristic (ROC) curves comparing OmniEM and EMDiffuse across all noise levels (one image montage; xy dimensions, 1596 × 1596 pixels). Scale bar: 200 nm.

**Extended Data Fig. 7.**
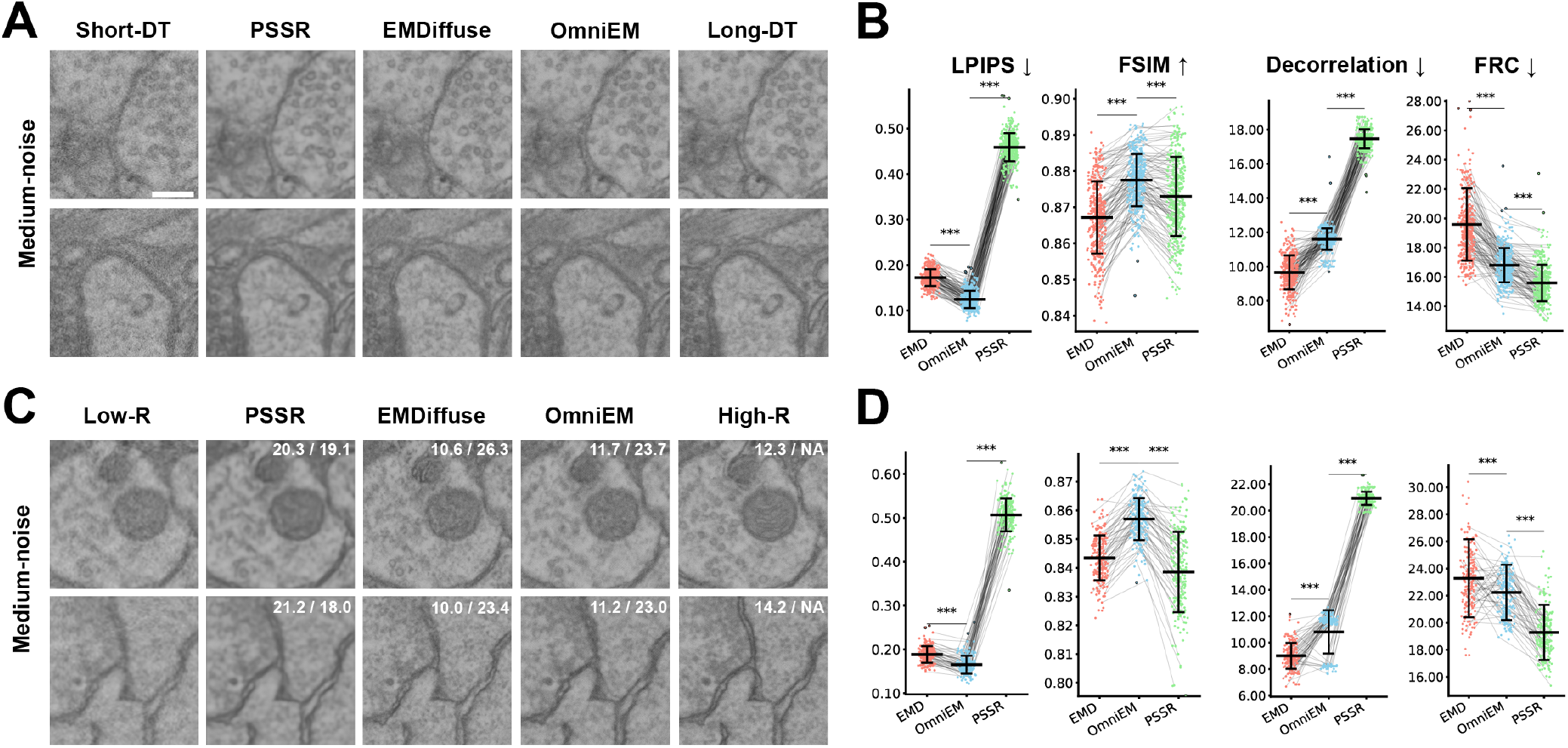
Medium-level denoising and super-resolution results. **(A)** Comparison of the denoising results from different models under medium-noise conditions. From left to right: the noisy image acquired with short dwell time (used as model input), the restored results from PSSR, EMDiffuse, and OmniEM, and the high-quality image acquired with long dwell time (used as ground truth reference). Two images are shown as representative cases. **(B)** Graphs comparing LPIPS, FSIM, and image resolution, assessed using decorrelation and FRC, across different models under medium-noise conditions. LPIPS, FSIM, and FRC resolution values were computed between each restored image and its corresponding ground truth, whereas decorrelation-based resolution was estimated independently of ground truth (n = 377 image tiles; xy dimensions, 224 × 224 pixels). Arrows next to each metric indicate the direction of better performance. Values are shown as mean ± s.d. P values are indicated for comparisons between OmniEM and the other two models (P < 0.05, P < 0.01, P < 0.001; NS, not significant; paired t-test). Lines connecting scatter points denote measurements from the same image tile restored by different models. For visualization clarity, 25% of data points are displayed. Data points exceeding mean ± 3 × s.d. were considered outliers, highlighted with black outlines, and excluded from statistical analysis. **(C)** Comparison of the super-resolution results from different models under medium-noise conditions. From left to right: the low-resolution input image, super-resolved outputs from PSSR, EMDiffuse, and OmniEM, and the high-resolution ground truth image. Two representative images are shown. Resolution values, estimated using decorrelation and FRC, are indicated in the upper right corner of each restored image. **(D)** Shown are the relative performance of different models on all pairwise test data under middle noise conditions for super-resolution, using the same four metrics. Layout and legend conventions follow those in B (n = 188 image tiles). Scale bar: 200 nm.

**Extended Data Fig. 8.**
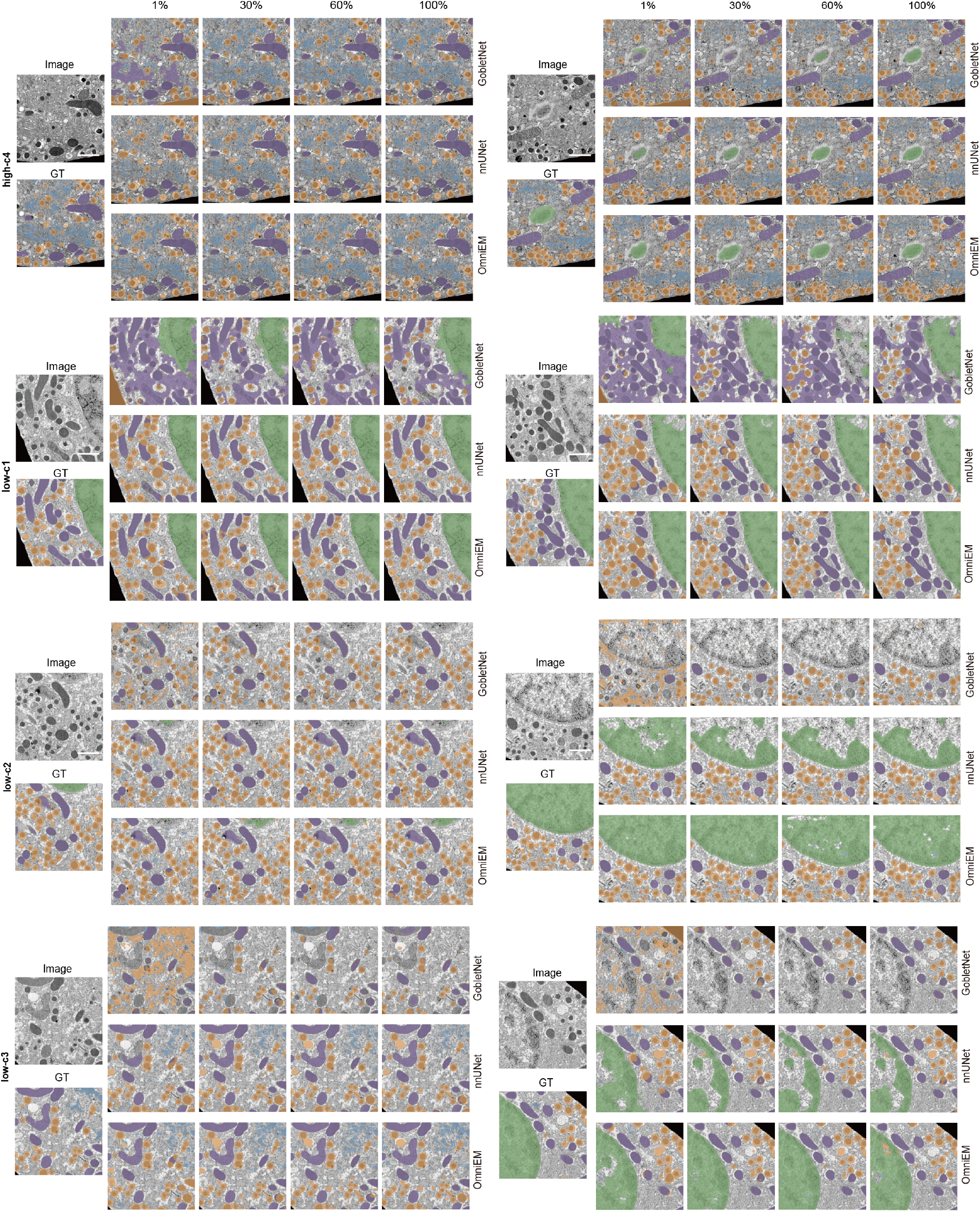
Multi-organelle segmentation results across BetaSeg EM volumes in 2D experiments. Example inference results on four BetaSeg volumes, shown from top to bottom: high-c4, low-c1, low-c2, and low-c3. The organization of sampled images follows the layout in Fig. 4C. Scale bar: 0.5*µ*m

**Extended Data Fig. 9.**
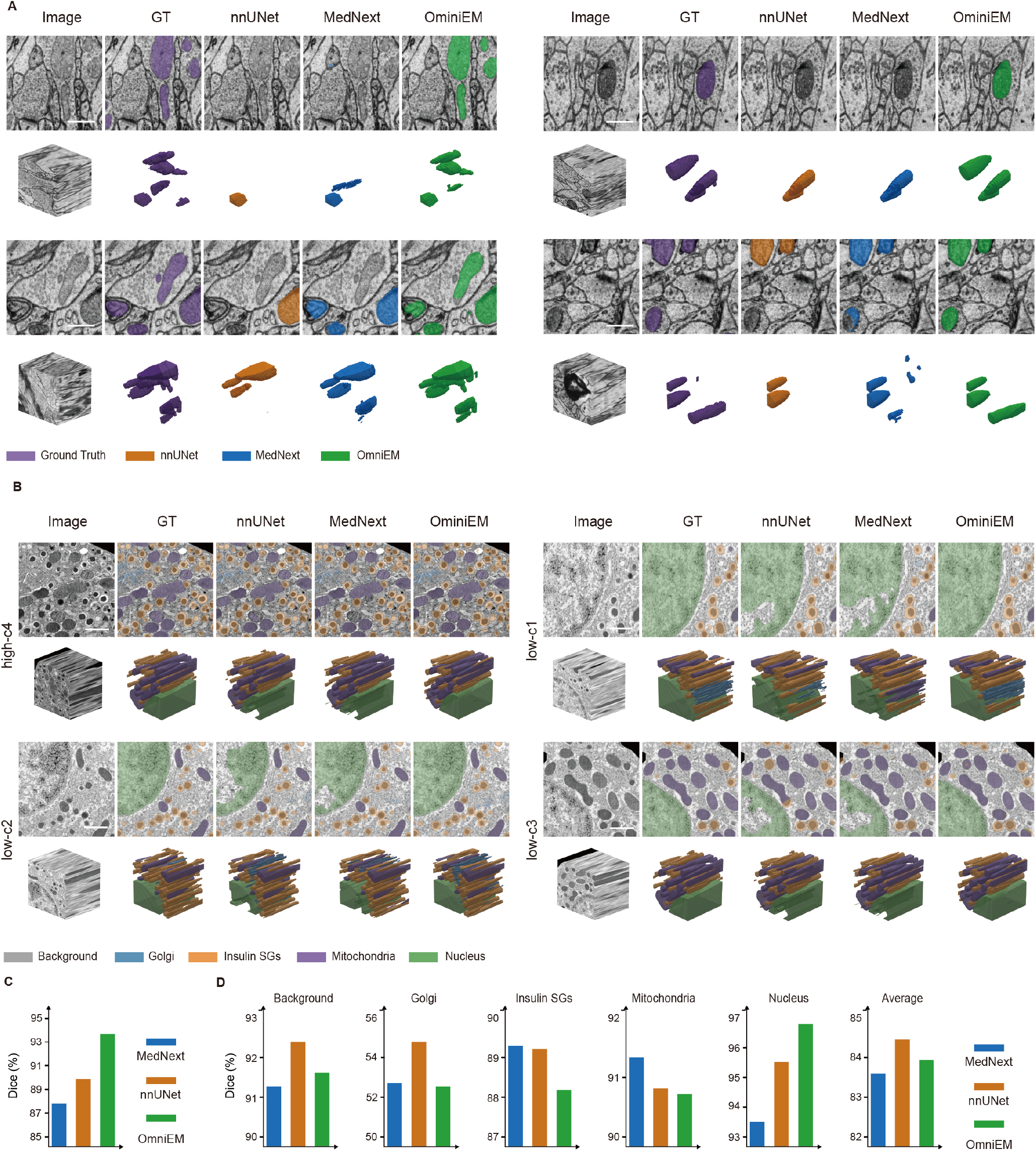
Visualization of 3D segmentation results. **(A)** Representative sub-volumes from the MitoEM-H dataset and the corresponding segmentation predictions. **(B)** Representative sub-volumes from BetaSeg (high_c4, low_c1, low_c2, low_c3, respectively) with their corresponding segmentation results, illustrating consistency across regions. Semantic legend definitions are the same as in Fig. 4D. **(C)** Mean dice coefficient comparisons between OmniEM (green) and MedNext (blue), and nnU-Net (brown) on MitoEM-H volume.**(D)** Mean dice coefficient comparisons between OmniEM (green) and MedNext (blue), and nnU-Net (brown) on four testing volumes from BetaSeg benchmark. Scale bar: 0.5*µ*m

## Footnotes

1 https://github.com/pku-maleilab/EM-SSL-project

2 https://histologyguide.com/EM-atlas/EM-atlas.html

3 https://histologyguide.com/EM-view/EM-133-cardiac-muscle/04-photo-1.html

4 https://histologyguide.com/EM-view/EM-347-megakaryocyte/08-photo-1.html

5 https://github.com/pku-maleilab/EM-SSL-project

## References

1. Loomba, S. et al. Connectomic comparison of mouse and human cortex. Science 377, eabo0924 (2022).

2. Xu, C. S. et al. An open-access volume electron microscopy atlas of whole cells and tissues. Nature 599, 147–151 (2021).

3. Witvliet, D. et al. Connectomes across development reveal principles of brain maturation. Nature 596, 257–261 (2021).

4. Svara, F. et al. Automated synapse-level reconstruction of neural circuits in the larval zebrafish brain. Nat. Methods 19, 1357–1366 (2022).

5. Parlakgül, G. et al. Regulation of liver subcellular architecture controls metabolic homeostasis. Nature 603, 736–742 (2022).

6. Shapson-Coe, A. et al. A petavoxel fragment of human cerebral cortex reconstructed at nanoscale resolution. Science 384, eadk4858 (2024).

7. Peddie, C. J. & Collinson, L. M. Exploring the third dimension: volume electron microscopy comes of age. Micron 61, 9–19 (2014).

8. Peddie, C. J. et al. Volume electron microscopy. Nat. Rev. Methods Primers 2, 51 (2022).

9. Collinson, L. M. et al. Volume em: a quiet revolution takes shape. Nat. methods 20, 777–782 (2023).

10. Kornfeld, J. & Denk, W. Progress and remaining challenges in high-throughput volume electron microscopy. Curr. opinion neurobiology 50, 261–267 (2018).

11. Buchholz, T.-O. et al. Content-aware image restoration for electron microscopy. Methods cell biology 152, 277–289 (2019).

12. Heinrich, L. et al. Whole-cell organelle segmentation in volume electron microscopy. Nature 599, 141–146 (2021).

13. Conrad, R. & Narayan, K. Instance segmentation of mitochondria in electron microscopy images with a generalist deep learning model trained on a diverse dataset. Cell Syst. 14, 58–71 (2023).

14. Aswath, A., Alsahaf, A., Giepmans, B. N. & Azzopardi, G. Segmentation in large-scale cellular electron microscopy with deep learning: A literature survey. Med. image analysis 102920 (2023).

15. Lu, C. et al. Diffusion-based deep learning method for augmenting ultrastructural imaging and volume electron microscopy. Nat. Commun. 15, 4677 (2024).

16. Popovych, S. et al. Petascale pipeline for precise alignment of images from serial section electron microscopy. Nat. communications 15, 289 (2024).

17. Lee, K., Zung, J., Li, P., Jain, V. & Seung, H. S. Superhuman accuracy on the snemi3d connectomics challenge. arXiv preprint arXiv:1706.00120 (2017).

18. Heinrich, L., Funke, J., Pape, C., Nunez-Iglesias, J. & Saalfeld, S. Synaptic cleft segmentation in non-isotropic volume electron microscopy of the complete drosophila brain. In International Conference on Medical Image Computing and Computer-Assisted Intervention, 317–325 (Springer, 2018).

19. Müller, A. et al. 3d fib-sem reconstruction of microtubule–organelle interaction in whole primary mouse β cells. J. Cell Biol. 220, e202010039 (2020).

20. Bae, J. A. et al. Functional connectomics spanning multiple areas of mouse visual cortex. Nature 640, 435–447, DOI: 10.1038/s41586-025-08790-w (2025).

21. Kievits, A. J., Lane, R., Carroll, E. C. & Hoogenboom, J. P. How innovations in methodology offer new prospects for volume electron microscopy. J. Microsc. 287, 114–137 (2022).

22. Dorkenwald, S. et al. Cave: Connectome annotation versioning engine. Nat. Methods DOI: 10.1038/s41592-024-02426-z (2025).

23. Awais, M. et al. Foundation models defining a new era in vision: a survey and outlook. IEEE Transactions on Pattern Analysis Mach. Intell. (2025).

24. Ma, C., Tan, W., He, R. & Yan, B. Pretraining a foundation model for generalizable fluorescence microscopy-based image restoration. Nat. Methods 1–10 (2024).

25. Sun, Y., Wang, L., Li, G., Lin, W. & Wang, L. A foundation model for enhancing magnetic resonance images and downstream segmentation, registration and diagnostic tasks. Nat. Biomed. Eng. 1–18 (2024).

26. Wang, X. et al. A pathology foundation model for cancer diagnosis and prognosis prediction. Nature 634, 970–978 (2024).

27. Archit, A. et al. Segment anything for microscopy. Nat. Methods 1–13 (2025).

28. Zhuo, Z. et al. Segment anything for dendrites from electron microscopy. arXiv preprint arXiv:2411.02562 (2024).

29. Wan, J. et al. Trisam: Tri-plane sam for zero-shot cortical blood vessel segmentation in vem images. arXiv preprint arXiv:2401.13961 (2024).

30. Iudin, A. et al. Empiar: the electron microscopy public image archive. Nucleic Acids Res. 51, D1503–D1511, DOI: 10.1093/nar/gkac1062 (2022).

31. Oquab, M. et al. Dinov2: Learning robust visual features without supervision. arXiv preprint arXiv:2304.07193 (2023).

32. Vo, H. V. et al. Automatic data curation for self-supervised learning: A clustering-based approach. arXiv preprint arXiv:2405.15613 (2024).

33. Fishbein, G. A. et al. Ultrastructural cardiac pathology: the wide (yet so very small) world of cardiac electron microscopy. Cardiovasc. Pathol. 73, 107670 (2024).

34. Schmitt, A., Guichard, J., Massé, J.-M., Debili, N. & Cramer, E. M. Of mice and men: Comparison of the ultrastructure of megakaryocytes and platelets. Exp. Hematol. 29, 1295–1302 (2001).

35. Fang, L. et al. Deep learning-based point-scanning super-resolution imaging. Nat. methods 18, 406–416 (2021).

36. Stites, H. C. & Manor, U. Pssr2: a user-friendly python package for democratizing deep learning-based point-scanning super-resolution microscopy. BMC Methods 2, 1 (2025).

37. Franco-Barranco, D. et al. Current progress and challenges in large-scale 3d mitochondria instance segmentation. IEEE transactions on medical imaging 42, 3956–3971 (2023).

38. Wei, D. et al. MitoEM dataset: Large-scale 3D mitochondria instance segmentation from EM images. In Lecture Notes in Computer Science, Lecture Notes in Computer Science, 66–76 (Springer International Publishing, Cham, 2020).

39. Roy, S. et al. MedNeXt: Transformer-driven scaling of ConvNets for medical image segmentation. (2023). 2303.09975.

40. Isensee, F., Jaeger, P. F., Kohl, S. A. A., Petersen, J. & Maier-Hein, K. H. nnU-Net: a self-configuring method for deep learning-based biomedical image segmentation. Nat. Methods 18, 203–211 (2021).

41. Zhou, Y., Li, L., Wang, C., Song, L. & Yang, G. Gobletnet: Wavelet-based high-frequency fusion network for semantic segmentation of electron microscopy images. IEEE Transactions on Med. Imaging (2024).

42. Le-Khac, P. H., Healy, G. & Smeaton, A. F. Contrastive representation learning: A framework and review. Ieee Access 8, 193907–193934 (2020).

43. He, K., Fan, H., Wu, Y., Xie, S. & Girshick, R. Momentum contrast for unsupervised visual representation learning. (2019). 1911.05722.

44. Chen, T., Kornblith, S., Norouzi, M. & Hinton, G. A simple framework for contrastive learning of visual representations. (2020). 2002.05709.

45. Conrad, R. & Narayan, K. Cem500k, a large-scale heterogeneous unlabeled cellular electron microscopy image dataset for deep learning. Elife 10, e65894 (2021).

46. Zhang, Y., Guo, J., Zhai, H., Liu, J. & Han, H. SegNeuron: 3D neuron instance segmentation in any EM volume with a generalist model. In Medical Image Computing and Computer Assisted Intervention – MICCAI 2024: 27th International Conference, Marrakesh, Morocco, October 6–10, 2024, Proceedings, Part VIII, 589–600 (Springer-Verlag, Berlin, Heidelberg, 2024).

47. Xie, R., Pang, K., Bader, G. D. & Wang, B. Maester: Masked autoencoder guided segmentation at pixel resolution for accurate, self-supervised subcellular structure recognition. In Proceedings of the IEEE/CVF Conference on Computer Vision and Pattern Recognition (CVPR), 3292–3301 (2023).

48. Li, Z. et al. Efficient masked autoencoders with self-consistency. (2023). 2302.14431.

49. Chen, R. J. et al. Towards a general-purpose foundation model for computational pathology. Nat. Med. 30, 850–862 (2024).

50. Wu, L., Zhuang, J. & Chen, H. VoCo: A simple-yet-effective volume contrastive learning framework for 3D medical image analysis. (2024). 2402.17300.

51. Popovych, S. et al. Petascale pipeline for precise alignment of images from serial section electron microscopy. Nat. Commun. 15, 289 (2024).

52. de Boer, P. et al. Large-scale electron microscopy database for human type 1 diabetes. Nat. Commun. 11, 2475 (2020).

53. Roels, J. et al. An overview of state-of-the-art image restoration in electron microscopy. J. microscopy 271, 239–254 (2018).

54. Pan, Y. et al. Adaptive template transformer for mitochondria segmentation in electron microscopy images. In 2023 IEEE/CVF International Conference on Computer Vision (ICCV), 21417–21427 (IEEE, 2023).

55. Shi, R. et al. ShapeMamba-EM: Fine-tuning foundation model with local shape descriptors and mamba blocks for 3D EM image segmentation. In Lecture Notes in Computer Science, Lecture Notes in Computer Science, 731–741 (Springer Nature Switzerland, Cham, 2024).

56. Lowe, D. G. Distinctive image features from scale-invariant keypoints. Int. journal computer vision 60, 91–110 (2004).

57. Motta, A. et al. Dense connectomic reconstruction in layer 4 of the somatosensory cortex. Science 366, eaay3134 (2019).

58. Xu, H. et al. A whole-slide foundation model for digital pathology from real-world data. Nature 630, 181–188 (2024).

59. McCafferty, C. L. et al. Integrating cellular electron microscopy with multimodal data to explore biology across space and time. Cell 187, 563–584 (2024).

60. Liu, Z. et al. Swin transformer: Hierarchical vision transformer using shifted windows. In Proceedings of the IEEE/CVF international conference on computer vision, 10012–10022 (2021).

61. Liu, X., Zhang, C. & Zhang, L. Vision mamba: A comprehensive survey and taxonomy. arXiv preprint arXiv:2405.04404 (2024).

62. Hu, E. J. et al. Lora: Low-rank adaptation of large language models. ICLR 1, 3 (2022).

63. Yu, J. et al. Unleashing the power of multi-task learning: A comprehensive survey spanning traditional, deep, and pretrained foundation model eras. arXiv preprint arXiv:2404.18961 (2024).

